# Untangling the effects of cellular composition on coexpression analysis

**DOI:** 10.1101/735951

**Authors:** Marjan Farahbod, Paul Pavlidis

## Abstract

**Background:** Coexpression analysis is one of the most widely used methods in genomics, with applications to inferring regulatory networks, predicting gene function, and interpretation of transcriptome profiling studies. Most studies use data collected from bulk tissue, where the effects of cellular composition present a potential confound. However, the impact of composition on coexpression analysis have not been studied in detail. Here we examine this issue for the case of human brain RNA analysis.

**Results:** We found that for most genes, differences in expression levels across cell types account for a large fraction of the variance of their measured RNA levels in brain (median R^2^ = 0.64). We then show that genes that have similar expression patterns across cell types will have correlated RNA levels in bulk tissue, due to the effect of variation in cellular composition. We demonstrate that much of the coexpression in the bulk tissue can be attributed to this effect. We further show how this composition-induced coexpression masks underlying intra-cell-type coexpression observed in single-cell data. Attempt to correct for composition yielded mixed results.

**Conclusions:** The dominant coexpression signal in brain can be attributed to cellular compositional effects, rather than intra-cell-type regulatory relationships, and this is likely to be true for other tissues. These results have important implications for the relevance and interpretation of coexpression in many applications.

## Background

Coexpression analysis is among the most-used methods in transcriptome data interpretation. The biological underpinnings of coexpression are well-established. Within a cell, genes whose products work together (either directly or indirectly) must be expressed together. This implies some commonality of regulation. Indeed, it is observed that genes with similar functions tend to be coexpressed (Eisen *et al.*, 1998; Langfelder *et al.*, 2011; Lee *et al.*, 2004; Gaiteri *et al.*, 2013). Based on these observations, coexpression is used in inferential frameworks (often via network-based approaches) to aid prediction of gene function and/or regulation (Li *et al.*, 2016; Rotival and Petretto, 2013; Amar *et al.*, 2013; de la Fuente, 2010; Saha *et al.*, 2017). In this paper, we examine assumptions that underlie such applications of coexpression to “bulk” samples of tissues containing multiple cell types. In particular, we explore the role played by variation in cellular composition.

In bulk brain tissue transcriptome datasets, gene expression clusters (sets of genes which are observed to be coexpressed) are often enriched for cell-type markers (Oldham *et al.*, 2008). Recently it has been proposed that variation in cell type composition between individual samples explains a substantial degree of variation in gene expression in human brain (Kelley *et al.*, 2018). In general, cell-type “deconvolution” methods rely on the idea that cell-type markers can be used to infer cellular composition (Newman *et al.*, 2015; Patrick *et al.*, 2019). Inferred cellular composition is also used for adjusting statistical models, as in some expression quantitative trait locus (eQTL) analyses (Westra *et al.*, 2015; Ng *et al.*, 2017). Thus there is at least implicit awareness that cellular composition is a factor in transcriptome data (Gaiteri *et al.*, 2013; Crow *et al.*, 2016). However, to our knowledge the connection between these observations and the interpretation of coexpression network analysis has not been described in detail.

In this study, we document the effect of cellular composition variability among samples in bulk nervous system tissue, and its downstream effect on network-based functional analysis. Using a combination of bulk tissue and single-cell data analysis, supplemented by simulations, we demonstrate that for a given gene the variance of its expression level in bulk tissue is directly related to its variability across cell types. We then show that this is strongly related to coexpression of genes with each other, such that the dominant signal in bulk tissue is simply due to variation in cellular composition across samples. Because many gene functions are highly associated with specific cell types, our results provide a major reason why clusters enriched for functions are observed in expression data. A further implication is that the utility of bulk tissue coexpression to infer transcriptional regulatory networks beyond uncovering cell-type specific expression patterns is greatly complicated. While our study focuses on expression in the human nervous system, the phenomena we document are likely to play an important role in analyses of other tissues.

## Results

### Variance of gene expression is highly affected by variation of cellular composition

Our work builds on two empirically-founded concepts. The first is that many genes are expressed at different levels in different cell types in the brain. The second is that brain tissue samples vary in their precise cellular composition. The latter occurs due to technical (e.g. sampling variability) and biological effects (von Bartheld *et al.*, 2016). The connection between the two in the context of bulk-tissue transcriptomics can be formalized in the following simple model, schematized in Figure 1 (for mathematical details see the Supplement). For each gene, we define a Cell Type (CT) expression profile, which is a vector of expression levels of the gene in each of *k* cell types. In the model, the CT profile is treated as a fixed intrinsic feature of the gene. Second, each bulk tissue sample has a specific cellular composition for those same *k* cell types. This forms a cellular composition vector of length *k* for each sample, where each element represents the proportion of a cell type in the sample. The observed expression level of a gene in the sample can be modeled as a weighted sum of the values in the CT profile, where the weights are given by the cellular composition vector of the sample. In the toy example shown in Figure 1, Gene 1 is only expressed in cell type B and therefore its relative expression in the data precisely tracks the proportion of cell type B present in the samples. This special case is used in many approaches to “cell type deconvolution”, where Gene 1 is considered a “marker gene” for cell type B. In contrast, Gene 5 is expressed equally in all cell types, so it is completely insensitive to differences in cellular composition and its expression level is the same in all the samples (for further mathematical details see Supplement section 1). The expression pattern becomes more complicated for a case like Gene 4, which is expressed at different levels in each of the two cell types, but because it is expressed at higher levels in cell type A, its pattern in the bulk tissue is positively correlated with the proportion of cell type A. Furthermore, genes that have correlated CT profiles will also be correlated in the bulk tissue (illustrated in Figure 1 by genes 1 and 2, and genes 3 and 4).

**Figure 1.**
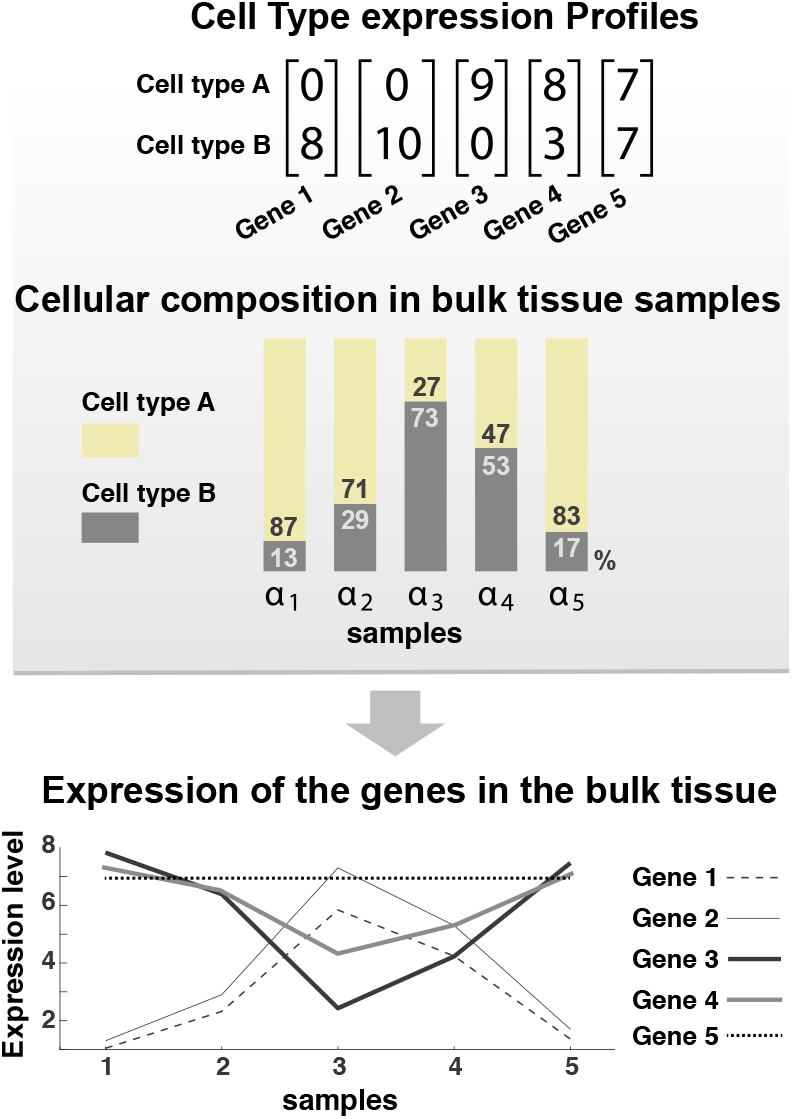
Schematic of cellular composition effects on gene expression variance in bulk tissue. **Top: Cell type (CT**) profiles for five genes in a hypothetical tissue with two cell types. Genes 1 and 2 are marker genes for cell type B. Gene 3 is a marker gene for cell type A. Gene 4 is expressed in both cell types, but at different levels, while Gene 5 is expressed at equal levels. Middle: Hypothetical cellular compositions of five bulk tissue samples. Each sample (***alpha***_**i**_) has the same amount of biological material but different proportions of each cell type. **Bottom:** The expected observed expression levels. Gene 1 and Gene 2 are positively correlated and negatively correlated with Gene 3 and 4. Gene 5 is expressed at the same level in all the bulk tissue samples as it is equally expressed in all cell types.

In general, the model predicts that the more variable the elements in gene’s CT profile, the more its measured expression in bulk tissue will be affected by variability in cellular composition of the samples (see supplement section 1 for simulation results demonstrating this). It is important to note that this model ignores all other potential sources of variability including noise or technical artifacts, as well as interactions between genes or cells that can influence expression. Our goal is to explore how well this effect explains the observed variance and correlation of genes in bulk tissue data.

As an initial assessment of whether this model is broadly explanatory, we estimated CT profiles for human cortex from single nucleus RNA-seq data (snuc-RNAseq data, see Methods), yielding expression levels for 16,789 genes in each of 75 different cell types, including all of the major classes of cells expected to be present in bulk cortex. We compared these data to a bulk cortex transcriptome dataset from GTEx (GTExBulk, see Methods). As predicted by the model, the variance of a gene’s expression in GTExBulk is correlated with the variance of its CT expression profiles (Spearman’s rho = 0.18; Supplementary Figure 3). Given the many potential sources of error, including noise in the CT profiles as well as the GTExBulk data, the agreement with the naive model is striking.

We next applied an approach related to many deconvolution methods to estimate the amount of variance attributable to cellular composition effects for each gene. As demonstrated in Figure 1, expression levels of cell-type marker genes in bulk tissue will reflect the variation of cellular composition among the samples. Therefore, the cellular composition-induced variance of the genes could be modeled by the variation of marker genes in a dataset. Here we used Principal Component Regression (PCR) using the expression of marker genes to predict the variation of the non-marker genes in the GTExBulk dataset (see Methods). The amount of variance explained by the model for each gene (R^2^) is an estimate of the degree to which the gene’s expression pattern is due to variability in cellular composition. In our first analysis, we used sets of marker genes from a high-quality snuc-RNAseq dataset for five major brain cell types (Pyramidal, Microglia, Astrocyte, Oligodendrocyte and Endothelial; see Methods). The resulting gene-level values of R^2^ range up to 0.91 (90^th^ quantile is 0.79) with a median of 0.46. In contrast, the same models fit to the snuc-RNAseq data, where we expect no effect of cellular composition (barring contamination of individual nuclei), the mean R^2^ is 0.018, with only 38 genes having values greater than 0.2.

As predicted by our model, R^2^ values are correlated with the variance of the CT expression profiles (rho = 0.28). To check the robustness of these findings, we tested another set of (largely non-overlapping) marker genes from Mancarci et al. (2017) with similar results (rho= 0.3; see Supplement section 2). We also tested randomly selected sets of non-marker genes instead of markers and found that R^2^ values are significantly higher for the marker genes when PCs are obtained from marker genes compared to the random selection of genes with similar average expression levels (p < 0.01 for average of R^2^ of marker genes for 100 trials, see supplementary Figure 02). Likewise, the two marker sets also generated higher R^2^ values for each other than the random gene sets despite their small overlap.

Motivated by reports that coexpression clusters are often associated with tissue-relevant gene functions, we next examined the relationship between gene function and cellular expression patterns. We observed that genes associated with brain-related functional terms (see Methods) tend to have higher R^2^ values, consistent with expected cell-type specific expression patterns in the brain (see Figure 2A). That is, genes with a brain-related function tend to have more varying CT profiles -- they are enriched in particular cell types -- which leads to high variation in bulk tissue. For example, genes involved in synaptic transmission are expressed in neurons, while genes involved in myelination are expressed in oligodendrocytes. Examples are genes annotated with “Regulation of synaptic plasticity” (GO:0048167, mean R^2^ = 0.76) and genes annotated with “Axon ensheathment” (GO:0008366, mean R^2^: 0.68; see Additional file 01). In contrast, terms for housekeeping functions tend to be associated with genes with lower R^2^ values (Examples: “Histone demethylation”, mean R^2^ value: 0.61 – GO:0016575; “spliceosomal snRNP assembly”, average R^2^ value: 0.53 – GO:0000387– see Additional File 02). In a closer examination, we also see that genes associated with the brain-specific term “Regulation of synaptic plasticity” have significantly higher variance in GTExBulk dataset compared to genes associated with the housekeeping term “Histone demethylation” (p = 0.005, ttest). In contrast, in the snuc-RNAseq cell population (a dataset expected to not have cellular composition effects) they have significantly lower variance (p = 4e-4; see Figure 2B). In summary, these results demonstrate that some of the observed variance of genes can be attributed to cell type composition variation, and this is especially true for genes with tissue-specific functions due to their tendency to also have cell-type specific expression patterns.

**Figure 2.**
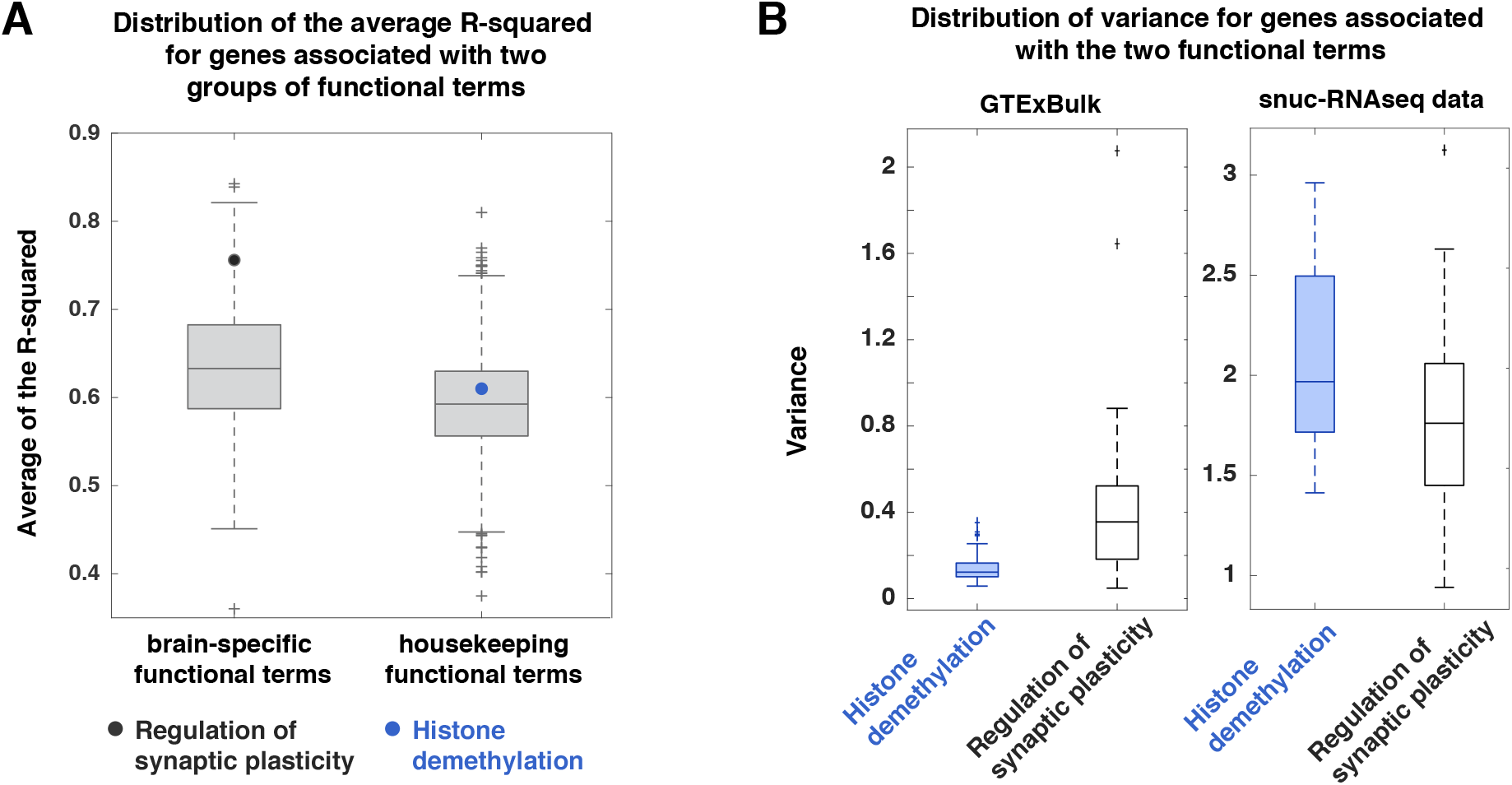
Much of the observed variance of brain-specific genes in bulk tissue is explained by cellular composition effects. **(A)** Groups of genes associated with brain-specific functional terms tend to have higher cellular composition-associated variance (indicated by higher average R^2^ values) compared to groups of genes associated with housekeeping terms. The average R^2^ values for two example terms are highlighted with black and blue dots. **(B)** Distribution of gene-level variance in the GTExBulk and snuc-RNAseq cell population (*Exc L2-3 LINC00507 FREM3*) for two groups of genes. Genes associated with brain-specific term “Regulation of synaptic plasticity” have higher variance than genes associated with the housekeeping term “Histone demethylation” in the GTExBulk dataset, while they have slightly but significantly lower variance in the sunc-RNAseq cell population.

### Much bulk tissue coexpression is explained by cellular composition variation among samples

In the previous section we demonstrated that variation in gene expression can be partly accounted for by variation in cellular composition. As illustrated in Figure 1, genes which have similar patterns of expression across cell types (as evidenced by correlated CT profiles) are also expected to have correlated expression in bulk tissue. Importantly, this phenomenon will be observed for any gene which has variability in expression across cell types, not just highly cell-type specific marker genes. For any two genes, in the absence of other factors, as the correlation between their CT expression profiles approaches one (or minus one), their correlation in bulk tissue is expected to approach one (or minus one – See supplementary section 1, Figure 1 and Figure 3C). We call this “cellular composition-induced coexpression”, to be distinguished from coexpression due to “within-cell” co-regulation. We hypothesized that it is a major source of observed coexpression in bulk tissue.

**Figure 3.**
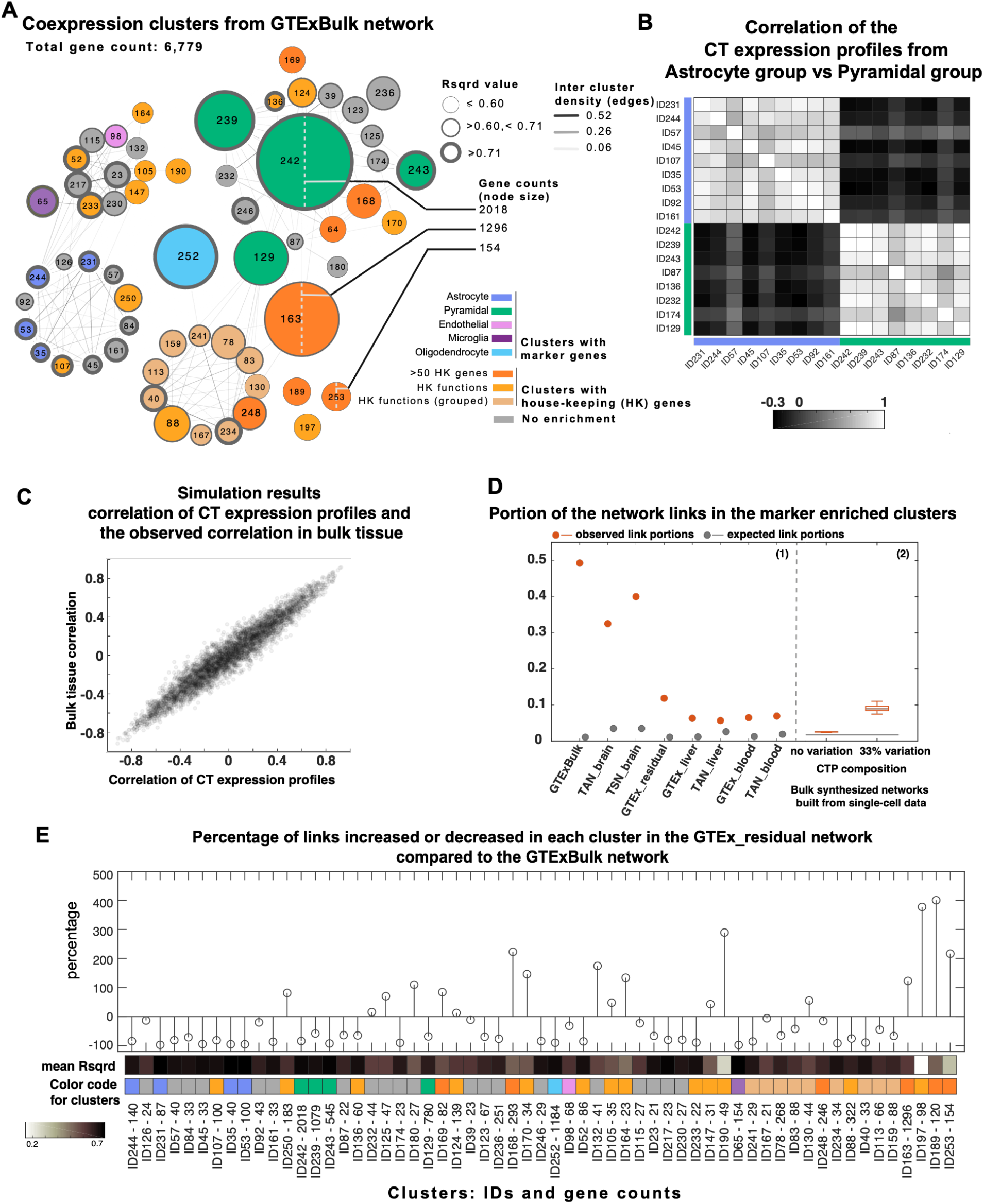
Much coexpression is explained by cellular composition effects. **(A)** Coexpression clusters from GTExBulk dataset network. Clusters are labeled with their IDs. Color indicates if the cluster has markers of a specific cell type, or if it is enriched with housekeeping functional terms or genes. Thickness of the border reflects the mean R^2^ value for genes in the cluster. **(B)** Coexpression of mean CT expression profiles for a group of clusters affiliated with Pyramidal cell expression patterns and a group of clusters affiliated with Astrocyte cell expression patterns (affiliation is indicated with presence of markers, high R^2^ or high inter cluster links) **(C)** Results from simulated bulk tissue data. Each dot represents data from a pair of genes. Plot shows data for 1000 gene pairs, sampled from a bulk tissue dataset with 100 samples and 10 hypothetical cell types. As demonstrated, for a given pair of genes, their Pearson correlation in the bulk tissue is highly correlated with the Pearson correlation between their CT expression profile. Also the higher the correlation between their CT expression profiles, the more likely their correlation in the bulk tissue is the same as the correlation of their CT expression profiles. **(D)** Proportion of coexpression involving the set of genes from the clusters enriched with marker genes. **(D1)** The two brain networks (GTExBulk and the TAN-brain) and the brain-specific network (TSN-brain) have between ~30-50% intra-cluster links in clusters enriched with marker genes. **(D2)** Portion of links in the same set of clusters in two groups of synthesized bulk tissue networks, modeling the effect of cellular composition variation. **(E)** Percentage of links increased or decreased for GTExBulk clusters in the residual network (GTEx_residual). Cluster color code as in A. The range for average R2 is shown in the color bar. Many housekeeping clusters (orange) with low average R2 values yield more links in the residual network.

We first performed clustering of the GTExBulk gene expression profiles, yielding 69 clusters (minimum 20 genes each; see Methods). As expected, some clusters are enriched with markers for one cell type (Figure 3A). While many of the other genes in these clusters are not markers, they tend to be “quasi-markers” – they are enriched in expression in a cell type (as for Genes 1,2 and 3,4 in the simplified model Figure 2 – see supplementary Figure 5 for expression levels of genes from clusters associated with different cell types, compared to marker genes). Furthermore, clusters that are enriched for markers for the same broad cell types and most of their neighboring clusters have correlated average CT expression profiles (0.993 > rho > 0.42, all p < 5e-4, Figure 3A and B). In addition, the average R^2^ values from the regression model are generally high for the marker-enriched clusters, consistent with composition-induced variance in expression (9/10 marker enriched clusters have average R^2^ greater than the median for all clusters (median = 0.65), Figure 3A; see Additional file 03 for all values).

The results so far make it apparent that some of the observed coexpression in the bulk brain tissue is explainable by cellular composition variation. Since cell-type specific patterns of expression are likely to be relatively fixed and therefore reproducible, composition-induced coexpression is also likely to be reproducible, and therefore contributing to the reported reproducibility of coexpression clusters among different bulk brain datasets. We examined this by comparing brain bulk tissue coexpression networks with each other and also with coexpression networks from other tissues. In the GTExBulk coexpression network, the intra cluster coexpression links in the clusters enriched with brain marker genes contain 49% of the total links. This is up to 40 times higher than the null expected value for the count of genes in the clusters for the given density of this network (see Figure 3D). The same set of genes have a high count of links in multi-dataset brain coexpression networks we previously described (Farahbod and Pavlidis, 2019), confirming that much of the reproducibility among bulk brain networks can be explained by cellular composition-induced coexpression. Importantly, while most (60-80%) of the genes in these clusters are also expressed in blood and liver, the high degree of observed coexpression among these particular genes is a phenomenon specific to the brain. We also see a large increase of links between the genes in marker enriched clusters in our simulated bulk tissue data upon the introduction of cellular composition variation (Figure 3D – see methods for details).

Apart from the marker-enriched clusters, many clusters in the GTEx-derived network are enriched with housekeeping genes and/or functions (see Figure 3). Most of these clusters have low mean R^2^ values (18/28 have mean R^2^ less than the median of all clusters (0.65) – see Additional files 03, 04), suggesting that their genes have small variability in their CT expression profiles and their coexpression is less likely to be affected by the cellular composition variation (like gene 5 in Figure 1). We hypothesized that some of the coexpression signal among genes from these clusters could have remained obscured due to the prevalence of high correlation values induced by cellular composition variation among other genes. To investigate this, we compared counts of links in different clusters in GTExBulk with counts of links in the GTExBulk residual network (GTEx_residual, a network built from the residuals of the PCR fits). We observed large drops in the count of links in the marker-enriched clusters and an increase in the count of links in clusters with low R^2^ values (Figure 3E). The magnitude of these changes highlight how cellular composition-induced coexpression can mask underlying coexpression within cell types. We discuss the use of the GTEx_residual network as a “corrected network” in the next section.

### Cellular composition effects can mask underlying intra-cell-type coexpression

We have shown that a major coexpression signal in bulk tissue comes from cellular composition effects. In our view this presents a shift from the usual interpretation, and raises the question of whether there is substantial coexpression attributable to other sources. This is especially relevant to attempts to infer coregulation. Specifically, the question remains as to whether coregulatory relationships in the sense typically sought are “visible” in bulk tissue data in the background of cellular composition effects. We do not attempt to fully address this question here, instead concern ourselves with a simpler one: in the common modes of coexpression analysis of bulk brain tissue, are coexpression patterns present *within* a cell type detectable? Composition-induced coexpression could in principle mask or amplify the bulk-tissue visibility of intra-cell-type coexpression. In this section, we examine the difference between robust intra-cell-type coexpression as measured in the snuc-RNAseq data and the observed coexpression in GTExBulk, and show that much of the difference can be attributed to the cellular composition effect. We also examine the GTEx_residual network as a form of “corrected network” in retrieving intra-cell-type coexpression.

As a preliminary step, we examined the general agreement of the coexpression networks built from different snuc-RNAseq populations and the GTExBulk dataset, and found that the agreement of network links is up to two times (and in few cases 3-5 times) higher for most of snuc-RNAseq populations than that expected by chance (see supplementary Figure 6). For reference, for our two bulk brain networks TAN-brain and GTExBulk this measure of agreement is 14. Conversely, some of the observed coexpression clusters in the GTExBulk are also reproducible in the larger snuc-RNAseq cell populations (see supplementary Figure 7), including those of many of the housekeeping and some of the brain-specific clusters. This shows that there is some level of agreement between coexpression observed in snuc-RNAseq and bulk data, in agreement with prior work (Crow *et al.*, 2016).

We then hypothesized that some of the differences between the observed coexpression in snuc-RNAseq data and GTEx bulk could be explained by the cellular composition effect, in a way that is shown schematically in Figure 4A. To test this, we compared bulk tissue coexpression with robust intra-cell-type coexpression patterns and found that for most part, differences between the two are explained by the effect of cellular composition variation. To identify robust intra-cell-type coexpression patterns, we combined 64 snuc-RNAseq coexpression networks built from neuronal cell types (both excitatory and inhibitory) and obtained a consensus (sum) “intra-cell-type” network (see Methods and Additional file 05 for list of links). We focused on the highest-confidence set of links that were present in 10 or more networks, leading to a set of 464 genes with 7,678 highly robust links. Of these links, 32% have correlation values above the 95^th^ quantile in the GTExBulk network but only 63% have correlation values above the median. This sub-network forms two very distinct clusters (Figure 4B). The gray cluster is enriched with multiple functional terms associated with neuronal processes and the black cluster is enriched with a few house-keeping functions (see the complete list of functions in Additional file 06). We then looked at the coexpression of these genes in the GTExBulk network (Figure 4C). While the genes in the two clusters have partly distinguishable coexpression patterns in GTExBulk, a large number of inter-cluster links are present; that is, the clusters are not as clearly separated. Indeed, the 464 genes appear in multiple clusters in the GTExBulk network (Figure 4D). We hypothesized that this is due to similarity of CT expression profiles among some of the genes, causing composition-induced coexpression that “blurs” the underlying cell-type-specific coexpression pattern. In support of this hypothesis, correlations of the CT profiles are high for some of the genes (Figure 4C and D). In particular, this can explain the differentiation between the two clusters ID253, ID168 and Excitatory cell clusters (indicated by green color bar in Figure 4D - these are the clusters enriched with Pyramidal markers – see Figure 3 for reference). The differentiation is even clearer when CT expression profiles are obtained from neuronal cells only (Figure 4E), indicating different expression patterns among neuronal cell types for genes in clusters ID253, ID168, ID64 and Excitatory clusters (Figure 4E). Our conclusion is that the intra-cell-type coexpression patterns observed in single cell data can be distorted and/or masked in bulk tissue by the effects of cellular composition.

**Figure 4.**
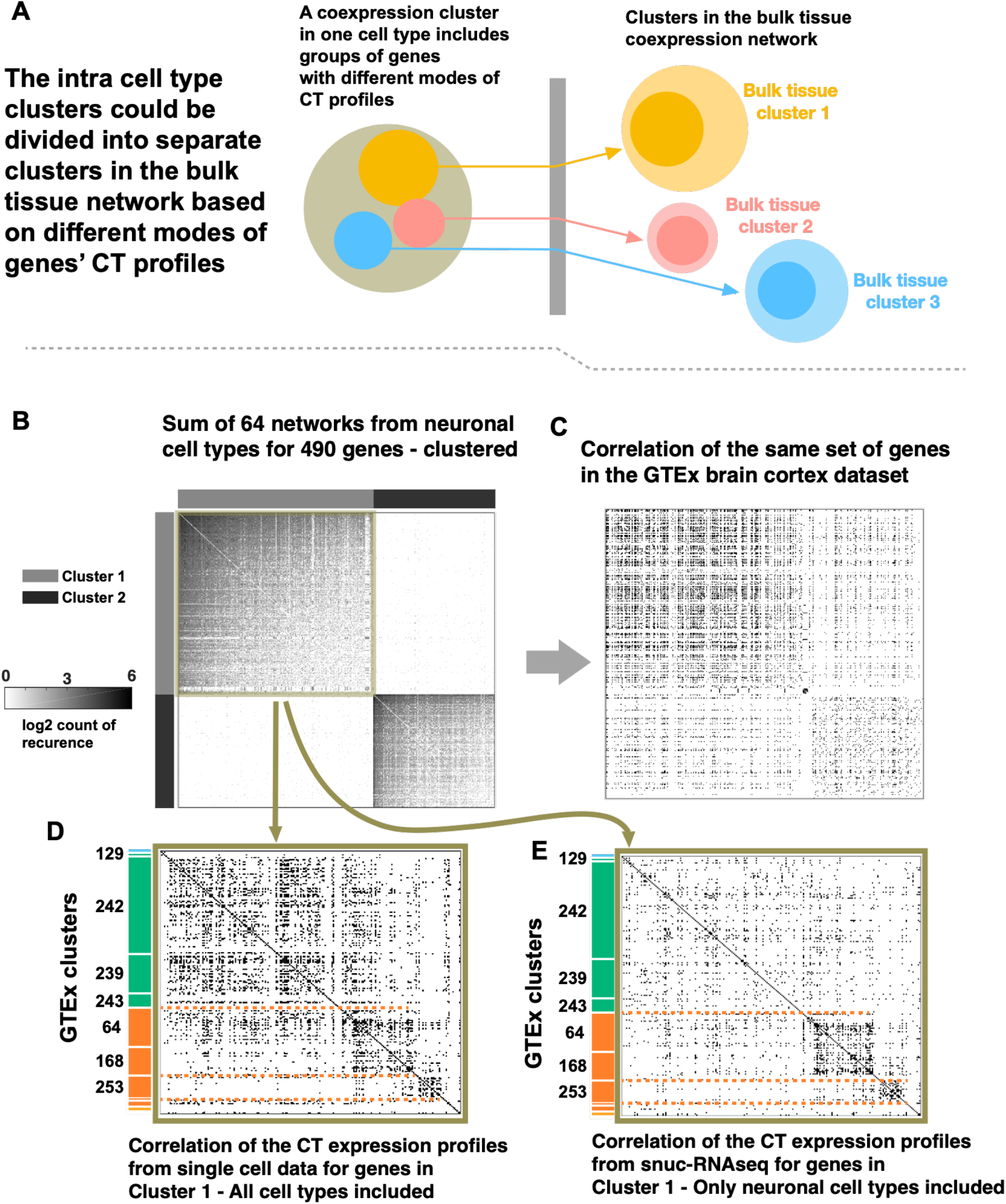
**(A)** Schematic showing a coexpression cluster in a specific cell type could be divided into multiple clusters in the bulk tissue dataset, as its genes might have different CT expression profiles. Each circle represents a group of genes. Colors blue, yellow and coral represent different “modes” of CT expression profiles, similar to the mean CT expression profiles for the bulk tissue clusters in Figure 3. **(B)** The heatmap shows part of the sum network from 64 neuronal snuc-RNAseq datasets where two coexpression clusters are identified. Clusters 1 and 2 (color bar grey and black) are well distinguished from each other. **(C)** Heat map shows the same set of genes with the same order in the GTExBulk tissue dataset. While the two clusters are somewhat distinguished, a great amount of intercluster links is present. **(D)** The heatmap shows the network for the genes in cluster 1, from the coexpression network built from the correlation of the CT expression profiles obtained from the 75 snuc-RNAseq datasets. Genes are ordered based on their belonging to different GTExBulk clusters, identified by the colorbar and cluster IDs from Figure 3A. Three sub-clusters are mildly distinguished, separating two groups of house-keeping clusters from the Pyramidal clusters (orange versus green). **(E)** Same plot as **D**, but the CT expression profiles are obtained from the 64 neuronal cell types only. The distinguished clusters demonstrate the group of genes with different expression levels in the neuronal cell types.

In the previous analysis, we showed that cellular composition effects can mask intra-cell-type coexpression especially when there is a conflict between the correlation of the CT expression profiles and the intra-cell-type coexpression, resulting in loss of the intra-cell-type pattern. In general, there are various scenarios that could occur, and intra-cell-type coexpression patterns might happen to be observed in bulk tissue to varying degrees and for varying reasons. Here we demonstrate this complexity with two genes, CALM3 and NRGN (Figure 5). They are robustly correlated in the snRNA-seq excitatory neurons (a link is present in 11 out of 23 of the networks for excitatory neurons), but there is no correlation between them in Inhibitory neurons, since NRGN is not expressed in Inhibitory neurons (Figure 5A, B). Accordingly, they have relatively highly correlated CT expression profiles (rho = 0.46), driven by their high expression in excitatory neurons and close to zero expression in the non-neuronal cell types, but moderated by their disjoint expression in inhibitory neurons. This suggests they might be coexpressed in bulk tissue, but for a reason different from that driving their coexpression within excitatory cells. As it happens, their correlation in bulk tissue is relatively high (97^th^ quantile), but not nearly high enough to pass our original 99.5 quantile filter for link selection. Their correlation ranks drop to the 82.3^th^ quantile in the GTEx_residual network. We conclude that the observed coexpression of CALM3 and NRGN in the bulk tissue is primarily caused by correlation of their cell type expression profiles, rather than a reflection of their coexpression in excitatory cells. Also, although their coexpression in the bulk tissue resembles their coexpression in excitatory cells, it is in disagreement with their lack of coexpression in other non-neuronal cell types. In general there is no simple relationship between coexpression within a cell type, and coexpression in a tissue in which that cell type is one of several present.

**Figure 5.**
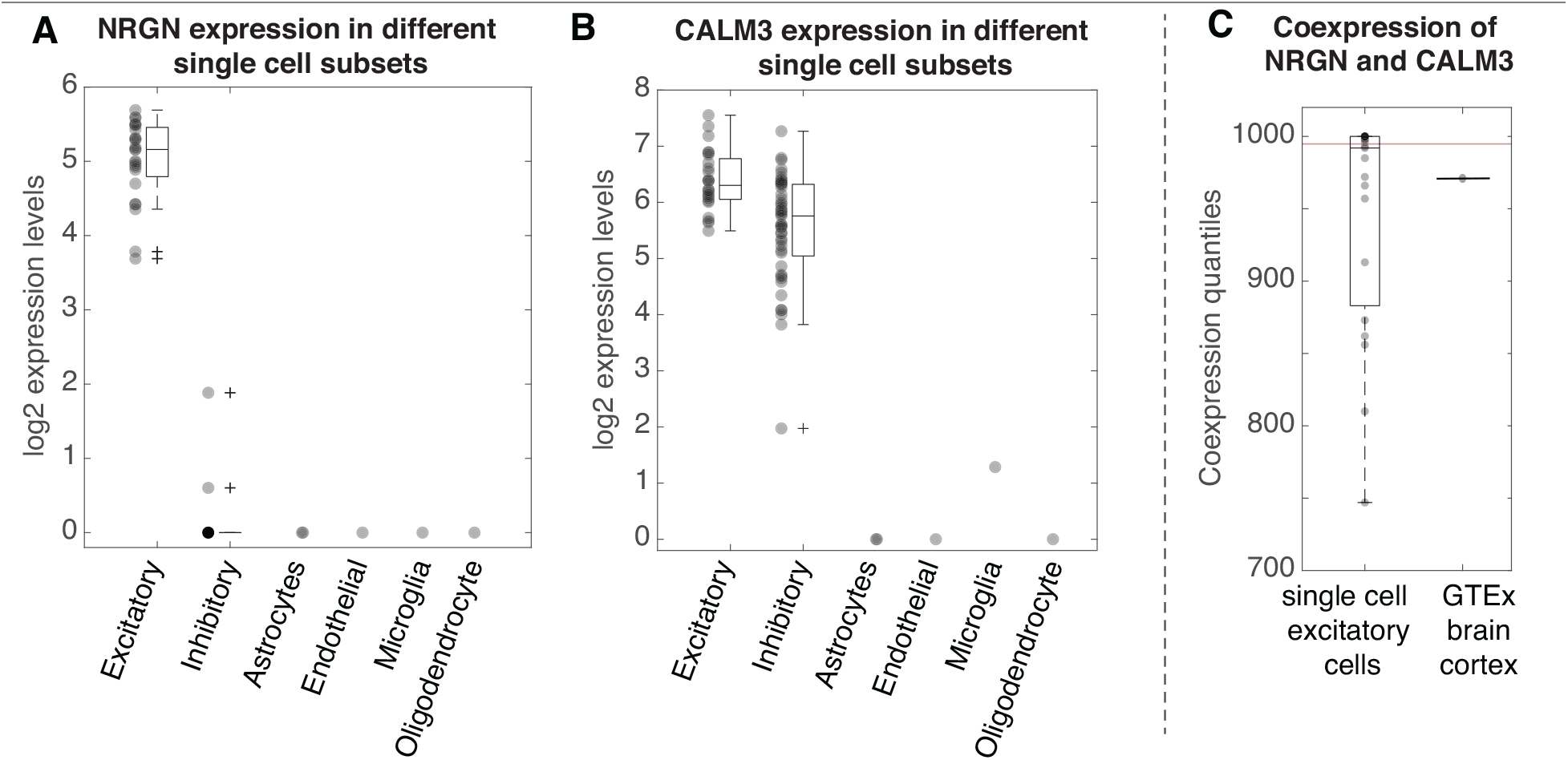
Coexpression of NRGN and CALM3 in Excitatory cell types. **(A)** From snuc-RNAseq data: NRGN is only expressed in the Excitatory cell types **(B)** From snuc-RNAseq data: CALM3 is expressed in both Inhibitory and Excitatory cell types **(C)** The two genes are highly correlated in the Excitatory cell types – based on coexpression networks built from snuc-RNAseq data. They are correlated in GTExBulk dataset but do not meet the threshold for the network (threshold is marked by the red line, it is the 995^th^ quantile).

Given that the effects of cellular composition on coexpression can be viewed as a confound, it is natural to consider whether the data can be corrected. In the previous section we observed that many of the clusters from GTExBulk had a much higher count of links in the residual network. In our framework a natural choice for such a correction are the residuals from our PCR model fits used to obtain the R^2^ estimates. We observe that most of the GTExBulk clusters are significantly reproduced in the GTEx_residual network (see supplementary Figure 6) and many of the brain-specific and housekeeping terms are enriched in the GTEx_residual network (Additional file 07). However, there is no overall significant improvement in agreement of the links in snuc-RNAseq populations with the GTexBulk_residual compared to the GTExBulk network (supplementary Figure 7), indicating that correction for composition may not be a panacea. But there is improvement of the precision in recovering snuc-RNAseq coexpression links for some of the GTExBulk-driven clusters. Results for the largest snuc-RNAseq population is presented in supplementary Figure 7. As an example we can consider the neuronal clusters with IDs 239, 242 and 243. These clusters are enriched for pyramidal cell markers, and their genes have high mean R^2^ values of 0.72, 0.74 and 0.75, respectively, suggesting that correction will have a large effect. In GTEx_residual the link density among the genes in these clusters decreases to approximately 1/2, 1/6 and 1/14 of the density in the GTExBulk network, while the link overlap with the snuc-RNAseq cell populations increases 24 to 175 percent. We also see that, for the most robust snuc-RNAseq clusters (from Figure 4B), which were substantially degraded in GTExBulk (Figure 4C), their separation is largely restored in the GTexBulk_residual network (Figure 6). However, the restoration is mostly due to the removal of inter-cluster links, rather than an increase in the intra-cluster cluster links. These findings suggest that correction for cellular composition effects can be beneficial, but confidence in the results are uncertain without matched single-cell data.

**Figure 6.**
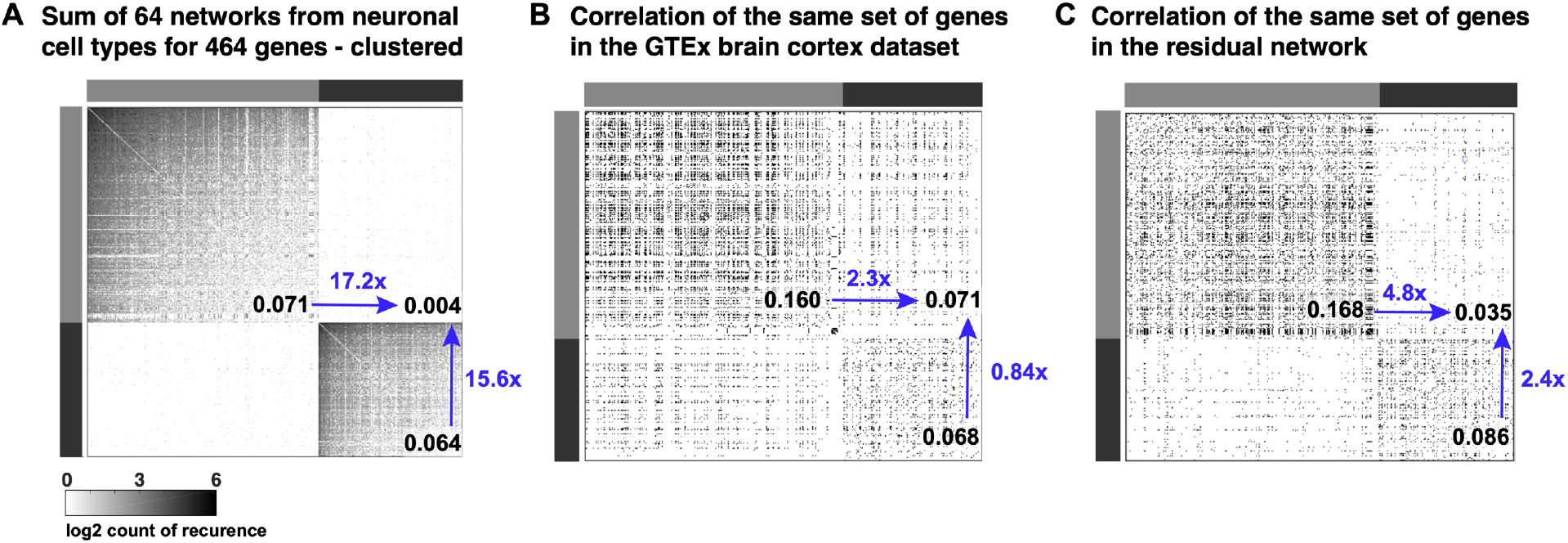
Reproducibility of robust correlation clusters from snuc-RNAseq networks in GTExBulk and the residual network. Inter and intra cluster density values are overlaid in black, and fold-changes in blue. The null density is 0.005 for all the networks **(A)** Sum network as in **Figure 4B**. For visualization, the data are scaled to be comparable to **B** and **C**. Intra cluster density is more than 15 times the inter cluster density for both black and gray clusters. **(B) As in Figure 4C**. Intra cluster density is less than the inter cluster density for the black cluster, for the gray one it is less than 2. **(C)** Intra cluster density is more than 10 times higher than the inter cluster density for both black and gray clusters. Notice that the dramatic effect in the density (from GTExBulk to GTEx_residual **network**) is mostly explained by the relative reduction of inter-cluster links rather than an increase in the intra-cluster links.

## Discussion

The term “coexpression” refers to two tightly linked concepts, one defined in the realm of molecular biology as the coordinated transcription of genes by regulatory mechanisms occurring within a cell (coregulation). The other is an observation of correlation of RNA levels in transcriptomic data; for clarity we can refer to the latter as “observed coexpression”. Coregulation has formed a foundation for understanding genome function for decades: that genes with collaborating products need to be expressed at the same time and are thus coregulated. Meanwhile, coexpression detected in high-throughput datasets has proven to be a reproducible signal with biological relevance: genes which are coexpressed have a higher probability of having related function than those which are not co-expressed. It is often assumed that observed coexpression is due to coregulation and therefore the latter could be reversed engineered from the former. Our work highlights a challenge, in that coexpression from bulk samples of a heterogeneous tissue are likely to be dominated by cellular composition effects. This has important implications for the interpretation of coexpression.

Coexpression in brain tissue is highly reproducible – that is, there are strong patterns of coexpression that are observed in many independent data sets (Farahbod and Pavlidis, 2019; Oldham *et al.*, 2008). Our results suggest that this is mostly due to the reproducibility of cell-type expression patterns, and the pervasive presence of variability in cellular composition between samples (Figure 1 and 3). We have shown that when coexpression is observed across cell types or tissues, the dominant patterns are due to cell-type or tissue-specificity of expression, and coexpression is merely a proxy for differential expression across cell types or tissues. While genes which are expressed specifically in one cell type (for example) can be thought of as having a “shared function”, that function is broad, only reflecting the function of that cell type. There is little expectation that function at the level of individual molecular interactions or pathways would be captured: the distinctness of a cell type cannot be fully described by the activity of a single pathway. Likewise, even for these genes their coregulation may reflect the broad epigenetic state of the cell type (Yoshida *et al.*, 2019), and finer-grained details of co-regulation are unlikely to be easily captured.

We have also shown that cellular composition-induced coexpression can mask apparently robust cell-type specific coexpression patterns (Figure 4). Despite this, a remaining question is whether correction for cellular composition would enable more efficient extraction of coregulation. For this to be the case, underlying patterns due to coregulation would have to be present in the data, and sufficiently separable from cellular proportion effects. For this to be effective, the regulation should ideally not be cell-type specific (otherwise the signal would be that much weaker; Figure 6), so the genes involved would have to be expressed in most cells. Since genes which are not cell-type specific tend to have housekeeping functions, it stands to reason that the most apparent coregulatory relationships would be those among housekeeping genes. We note that schemes for correcting bulk tissue data for cell type proportions (either directly or indirectly) are often used in expression QTL studies, and have been shown to increase the number of cis-eQTLs that can be recovered (Ng *et al.*, 2017). This suggests that correcting for cell type proportions and recovering underlying biological signals is possible, but eQTL studies require large sample sizes (generally at least 100 but often far more, especially for trans-eQTLs). We expect that identification of coregulatory relationships from bulk tissue data will similarly require very large samples sizes and still be most effective at extracting regulation of housekeeping genes rather than cell-type specific genes. Giving these constraints, it would seem preferable to use coexpression data from a single cell type to extract regulatory relations. However, limitations of the most commonly-used single-cell transcriptome methods suggest extracting high-quality regulation information is a challenge (Crow and Gillis, 2018). Furthermore, the most commonly used computational method for doing so is designed to simultaneously identify cell types along with building a regulation network, so that the strongest patterns observed are likely dominated by differential expression across cell types, not coexpression within cell types (Aibar *et al.*, 2017).

Our study does have some limitations. First, our analysis of cell-type-level coexpression is based on a single (albeit large and unusually deeply sequenced) data set that used different samples than the bulk tissue. Thus we cannot rule out that the failure to recover some snuc-RNAseq coexpression patterns in bulk tissue might reflect data-specific effects. This might be resolved in the future with additional data sets. Second, we only considered the phenomenon in brain. Intuitively, cellular composition effects should impact any bulk tissue coexpression analysis, but determining whether the inferred effects in other tissues are weaker or stronger than those we observe for brain should be a topic of future research. Finally, the actual cellular composition of the bulk tissue samples we used is not known. While the approach of using cell-type markers to infer composition has been validated many times (Newman *et al.*, 2015; Patrick *et al.*, 2019; Mancarci *et al.*, 2017), we do not claim it is a perfect substitute for accurate direct counts. It remains formally possible that some of the variation we attribute to cellular composition is instead due to complex patterns of gene regulation that mimic compositional effects, but we feel the most parsimonious interpretation of the data is that cellular composition is a major contributor. It is also worth noting that imperfect cell-type-effect measurement could just as easily cause us to underestimate the impact of composition, as the residual would still contain compositional effects.

Beyond the implications for the goal of inferring regulation, our results have important implications for any use of expression data-based gene clustering or module identification in which the patterns are driven by cellular composition effects. First, the representation of the data as a network is potentially misleading, because it is tempting to interpret a network as representing physical relationships. In particular, the idea that “hubs” in coexpression models are especially interesting is highly questionable if that pattern is simply a reflection of the cellular distribution of those transcripts. Second, if cellular composition is of interest, it would be reasonable to analyze composition more directly by inspecting the expression of known markers rather than by using indirect means via clustering and enrichment analysis. This parallels the situation for analysis of differential expression, where changes in measured expression levels can be due to changes in composition (Mancarci *et al.*, 2017; Toker *et al.*, 2018). On the other hand, machine learning applications of coexpression to tasks such as gene function prediction are not directly affected by our findings, as success in prediction does not necessarily depend on the biological meaning of the features used.

## Conclusions

For more than two decades, coexpression analysis has been among the most widely applied methods in genomics. Its popularity is based on the assumption that it is a window into gene regulation. Here we have shown that coexpression in bulk tissue could better be described as providing a window into the distribution of transcripts with respect to cell types. While this is useful information, this shift in interpretation should be considered in future studies. Coexpression remains an interesting phenomenon worthy of study, and our work contributes to greater understanding of its meaning and limitations, and we hope it leads to more informed data analyses.

## Methods

### Data

We have three main sources: 1. A single-nucleus dataset from

Allen Brain Atlas from Middle Temporal Gyrus (© [2018] Allen Institute for Brain Science. Cell Diversity in the Human Cortex. Available from: [http://celltypes.brain-map.org/download#transcriptomics]) 2. GTEx RNA-seq expression dataset from brain-cortex (Lonsdale *et al.*, 2013). 3. A set of coexpression networks: binary coexpression networks were built from GTEx RNA-seq blood and liver and a set of Tissue Aggregated Networks (TANs) from blood, brain and liver, from our previous study (Farahbod and Pavlidis, 2019). The TAN networks are built by aggregating several networks from each tissue, built from datasets on Affymetrix platform. The TSN-brain network is a subset of the TAN-brain network, where the links are identified as specific to the brain, blood or liver. Supplementary table 1 provides counts of genes and links in each of the networks. All data and scripts used for the analysis are available from the authors.

### Single-nucleus data

The snuc-RNAseq dataset has records from 15,928 nuclei for a total of 50,281 genes, grouped into 75 cell types. We used the read counts from exons only and did not use the intronic reads. We used the labels for the cell-types based on the clustering provided by the Allen Institute. We removed nuclei which had data for less than 2000 genes and nuclei for which the total read count was more than 3 times or less than 1/3 times the median. Genes were filtered for NeuN negative and NeuN positive samples separately. We selected genes expressed in at least 2% of the nuclei or expressed at the highest quartile in the nuclei it is expressed in. The final dataset has data for 16,789 genes and 15,646 nuclei. Supplementary Figure 2 shows the count of cells in each group of the 75 cell types. To construct Cell Type (CT) vectors for each gene, we obtained the mean expression level of that gene in each of the 75 cell types. Each of the 16,789 genes yields a CT expression profile vector of 75 elements.

We built coexpression networks for 69 of the 75 cell-types in the snuc-RNAseq data set (six cell-types had too few cells). For each cell type, correlations were computed for each pair of genes using only nuclei in which expression was greater than zero to reduce the impact of drop-outs. Gene pairs with less than 20 usable nuclei were removed. Because of differences in sample size for the correlations (causing different null distributions for the correlation), we omitted the correlation threshold filtering step used for the other data sets, and therefore filtered the one-sided p-values of the correlations (METHOD FOR PVALUES) to identify the 0.5% of the gene pairs with smallest p-values. Supplementary Table 2 has the link count and gene count for the 69 networks.

To construct combined networks, we summed the 64 binary coexpression networks built from Inhibitory and Excitatory neurons. Robust coexpression links were identified as those present in 10 or more of the networks, between genes with more than two such links. The total of 490 genes passed this criterion and were clustered using topological overlap and hierarchical clustering. Sixteen mitochondrial genes and 10 unclustered genes were removed (the presence of mitochondrial genes is likely due to variable mitochondrial contamination of the nuclei). The remaining grey and black clusters have 286 and 178 genes respectively.

### GTEx datasets and networks

The read counts per million reads (CPM) values from each of the three GTEx datasets: brain-cortex, liver and blood were filtered to include the genes with CPM > 0 in > 20% of the samples. Expression values were log_2_ transformed and binary coexpression networks were built using the Pearson correlation, filtered to include the 0.5% of the links with highest correlation values in each of the three networks. The counts of links and genes included in each network are provided in Supplementary Table 1. To cluster the GTExBulk network, we applied hierarchical clustering to the Topological Overlap (TOP) (Zhang and Horvath, 2005). An initial set of 253 clusters were identified for 12,416 of genes, of which 69 clusters had at least 20 genes and were retained for further study. Clustering labels are in Additional file 0.8. Counts of marker genes for different cell types for each cluster are in supplementary Figure 2.

### Identification of marker genes

Marker genes were selected based on two sources. From snuc-RNAseq data, for each of the five major cell types Astrocyte, Oligodendrocyte, Microglia, Endothelial and Pyramidal (labeled as Excitatory cell types), we identified genes with mean count per million ≥ 2 fold-change in all other cell-types as marker genes. Additional file 09 has list of markers identified by this manner for each of the five cell types.

As the second source, we used 1,208 marker genes for 18 mouse cerebral cortex cell types identified by Mancarci et al. (2017). We mapped the the marker genes to their human orthologs using the Ensembl database (Zerbino *et al.*, 2018). Mancarci et al. further refined these markers based on their coexpression in bulk human tissue. To remain consistent with their method, for each cell type we only considered the subset of its marker genes which were highly correlated with each other, using the hierarchical clustering with topological overlap on the marker genes for each cell type. Genes in the cluster with highest count of links were selected as the markers of the cell type. Our final list includes 256 markers for five major cell types (Additional file 10). Supplementary figure 2 shows overlap of the marker gene sets from the two sources.

### Modelling expression level of genes in the GTExBulk dataset

We used linear models to estimate the expression level of genes based on the variation of the marker genes in each of the samples, using the first seven principal components of the whole set of marker genes (the snuc-RNAseq and Mancarci (2017) marker sets separately) in the bulk tissue dataset. Therefore, the expression level of gene *A* in sample *j* of the GTExBulk dataset is modeled as:

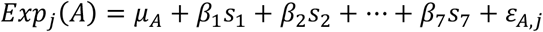

Where μ_A_ is the average expression level of gene *A*, *s_i_*’s are the principal component scores for sample *j*, β_j_’s are the parameters of the model and ε_A,j_ is residual error.

### Enrichment of functional terms

For a given network, each functional term is marked as enriched if the density of the links between the genes associated with the term are significantly higher than the density of the network, where the hypergeometric distribution is used as the null. The FDR was controlled at 0.1 using the method of Benjamini and Hochberg (Benjamini and Hochberg, 1995). Brain-specific functional terms were identified as the terms enriched in either of the TAN-brain networks for GTExBulk network, but not enriched in TAN-liver and TAN-blood networks. Likewise, housekeeping terms were identified as terms enriched in either of the TAN-brain or GTExBulk networks, as well as the TAN-blood and TAN-liver networks.

### Synthesized bulk datasets

Each of synthesized sample was built using snuc-RNAseq samples (nuclei) from the Allen data described above, as follows. First, nuclei were grouped into five major cell types based on their provided labels. Then, for each synthetic sample, nuclei were randomly sampled with the following baseline proportions: Pyramidal (20%), Inhibitory (20%), Oligodendrocyte (43%), Astrocyte (12%) and microglia (5%), based on the estimates of Von Bartheld et al. (2016), and their gene expression values were added and divided by the total count of nuclei, yielding a final synthetic sample. This was repeated to create multiple synthetic data sets with 100 samples each. To add composition variability, the baseline proportions were randomly varied by drawing each proportion from a normal distribution with variance 33% of its mean.

## Supporting information

Functional terms enriched in brain network

Summary information for GTEx clusters

Robust snuc-RNAseq links

Clustering label and R2 values for genes in GTExBulk

functional terms enriched in brain and at least one other tissue (common terms)

Functional enrichment results for GTExBulk and GTEx_residual networks

Functional terms enriched in GTExBulk clusters

Final list of marker genes from Mancarci et al.

List of cell type markers from snuc-RNAseq

## Declarations

## Availability of data and material

The GTEx dataset analyzed in this study is available from GTEx portal at [https://gtexportal.org/home/datasets] doi: 10.1038/ng.2653.

The single-nucleus dataset Cell Diversity in the Human Cortex, from Allen Institute for Brain Science, analyzed in this study is available at [http://celltypes.brain-map.org/download#transcriptomics]

The Tissue Aggregated Networks (TANs) used in this study are available in the supplementary material of the article “Differential coexpression in human tissues and the confounding effect of mean expression levels” doi: 10.1093/bioinformatics/bty538.

## Authors’ contributions

PP and MF conceived the study. MF conducted research. PP provided oversight. MF and PP wrote the manuscript.

## Acknowledgements

We thank the Allen Institute for Brain Science and the GTEx consortium for making their data available, without which this study could not have been conducted. We thank Shreejoy Tripathy, Ogan Mancarci and members of the Pavlidis lab for advice and discussion.

## Additional files

- Additional file 1
- .tsv
- functional terms enriched in brain network
- The file contains list of Gene Ontology terms that are identified as brain-specific. It has four columns: GOID [Gene Ontology ID], GOTerm [Gene Ontology term], Gene Count [count of genes associated with that term], MeanRsqrd [mean Rsquared value of the genes associated with the term]

- Additional file 2
- .tsv
- functional terms enriched in brain and at least one other tissue (common terms)
- The file contains list of Gene Ontology terms that are identified as brain-specific. It has four columns: GOID [Gene Ontology ID], GOTerm [Gene Ontology term], Gene Count [count of genes associated with that term], MeanRsqrd [mean Rsquared value of the genes associated with the term]

- Additional file 3
- .txt
- Summary information for GTExBulk clusters
- The file has three columns: ClusterID [ID of the cluster], memberCount [count of genes in the cluster], averageR2 [average of Rsquared value for genes in the cluster]

- Additional file 4
- .txt
- Functional terms enriched in GTExBulk clusters
- For each cluster ID, enriched functional terms are written. Only terms with FDR ≥ 0.1 were included. The file has three columns: GOID [Gene Ontology ID for the enriched term], GOTerm [the enriched term], pvalue [pvalue of the enrichment].

- Additional file 5
- .txt
- Robust snuc-RNAseq links
- Each row has information about one of the links and the file has nine columns: gene_1 [gene symbol of one of the genes], gene_2 [gene symbol of the other gene], totalRep [total count of snuc-RNAseq networks that the link is present in], inhRep [count of Inhibitory neuron networks that the link is present in], excRep [count of Excitatory neuron networks that the link is present in], GTExCorrRank [rank of the link in GTExBulk network, 1000 is the highest correlation values and 0 is lowest], TAN_presence [the link is present in TAN-brain network or not], gene1_r2 [Rsquared value for gene_1], gene2_r2 [Rsquared value for gene_2]

- Additional file 6
- .txt
- Functional terms enriched in robust snuc-RNAseq cluster – the gray cluster
- The file has three columns: GOID [Gene Ontology ID], GOTerm [Gene Ontology term], FDR [FDR from the enrichment]

- Additional file 7
- .txt
- Functional enrichment results for GTExBulk and GTEx_residual networks
- The file has four columns: GOID [Gene Ontology ID], GOTerm [Gene Ontology term], GTExBulk_p [corrected p value for GTExBulk network], GTEx_residual_p [corrected p value for GTEx_residual network]

- Additional file 8
- .txt
- Clustering label and R2 values for genes in GTExBulk
- The file has three columns: GeneSymbol, clusterID, R2

- Additional file 9
- .txt
- List of cell type markers from snuc-RNAseq
- Each column is labeled with a cell type and contains list of marker genes obtained from snuc-RNAseq data

- Additional file 10
- .txt
- Final list of marker genes from Mancarci et al.
- Each column is labeled with a cell type and contains the final list of markers obtained from Mancarci et al.

