## Supplementary file for "Untangling the effects of cellular composition on coexpression analysis"

### Contents

1 **Modeling and Simulation** **1**

1.1 Simulation 1 - variance of a gene in bulk tissue . . . . . . . . . . . . . . . . . . . . 2

1.2 Simulation 2 - correlation of two genes in bulk tissue . . . . . . . . . . . . . . . . . 3

2 **Gene variance explained by variance of the marker genes** **4**

3 **Figures** **5**

4 **Tables** **13**

### 1 Modeling and Simulation

For a given gene $q$ and a tissue with $m$ cell types, we use the vector $\mathbf{c}_q$ to represent its Cell Type expression profile (CT profile), which contains the expression level of $q$ in each of the $m$ cell types:

$$\mathbf{c}_q = [\,t_{q1},\, t_{q2},\, ...,\, t_{qm}\,], \qquad t_{q1}, t_{q2}, ..., t_{qm} \geq 0,\, \sum_{i=1}^{m} t_{qi} > 0 \tag{1}$$

Where $t_{qi}$ is the mean expression level of $q$ in cell type $i$. We denote the variance of values $t_{qi}$s by $var(t_q)$, that is, the variance of the expression level of $q$ among different cell types.

For a given dataset with $n$ samples from the tissue, vector $\boldsymbol{\alpha}_j$ represents the Cell Type Composition (CTC) in sample $j$:

$$\boldsymbol{\alpha}_j = [\,\alpha_{j,1},\, \alpha_{j,2},\, ...,\, \alpha_{j,m}\,], \qquad \alpha_{j,i} \geq 0, \sum_{i=1}^{m} \alpha_{j,i} = 1 \tag{2}$$

Where $m$ is the count of cell types in the tissue, as it was in definition 1 and $\alpha_{j,i}$ is the proportion of cell type $i$ in sample $j$. Matrix $\mathbf{A}$ defined as:

$$\mathbf{A} = \left[\boldsymbol{\alpha}'_1,\, \boldsymbol{\alpha}'_2\,, ..., \boldsymbol{\alpha}'_n\right] \tag{3}$$

contains cellular composition vectors for the $n$ samples of the dataset.

##### 1.1 Simulation 1 - variance of a gene in bulk tissue

Having the above definitions, the expression level of $q$ in $n$ samples of the dataset is given as $\mathbf{e}_q$:

$$\mathbf{e}_q = \mathbf{c}_q\,\mathbf{A} \tag{4}$$

We denote the variance of the expression level of $q$ among the $n$ samples of dataset presented in $\mathbf{e}_q$ by $var(q)$. From the above formulation, it is apparent that for the two special cases when $t_{q1} = t_{q2} = ... = t_{qm}$, (i.e $var(t_q = 0)$ or when $\boldsymbol{\alpha}_1 = ... = \boldsymbol{\alpha}_n$, we have $var(q) = 0$. In a bulk tissue dataset, these two cases are equivalent to when $q$ has the same expression level in all cell types and when the cellular composition remains the same among the samples.

For simulation, vectors $c$ (CT expression profiles) were generated for $1000$ genes for $m = 10$ cell types. For each gene $q$, $\mathbf{c}_q$ was generated using normal distribution with mean and variance obtained from a CT expression profile of a randomly selected gene from the snuc-RNAseq dataset. Bulk tissue data was generated with $n = 100$ samples. Matrix $A$ was generated using uniform distribution with the criteria that each column has sum one. The results show that among the genes, $var(t_q)$ and $var(q)$ are highly correlated (see supplementary Figure 03).

###### MATLAB code for simulation

---

```
myVar % variance of the CT expression profiles obtained from snuc-RNAseq data
myMean % mean of the CT expression profiles obtained from snuc-RNAseq data

m = 10 % count of tissues
n = 100 % count of bulk samples

% normalizing the weight matrix
A = rand(m,  n);
for i = 1:n
    A(:, i) = A(:, i) ./(sum(A(:,i)));
end

orVar = zeros(1,1000); % variance of the CT profile
obVar = zeros(1,1000); % the observed variance in bulk tissue
```

```
for k = 1:1000 % for 1000 genes
    ind = datasample(1:length(myVar), 1);
    v = myVar(ind);
    orVar(k) = v;
    c = normrnd(myMeans(ind), sqrt(v), 1, t);
    exps = c * A;
    obVar(k) = var(exps);
end
```

---

##### 1.2 Simulation 2 - correlation of two genes in bulk tissue

For given two genes $p$ and $q$, with expression vectors $\mathbf{e}_p$ and $\mathbf{e}_q$ in the bulk tissue dataset, the higher the correlation of their CT profiles $\mathbf{c}_p$ and $\mathbf{c}_q$ (in any direction), the more likely that $\mathbf{e}_p$ and $\mathbf{e}_q$ are also correlated in the same direction. Having centered $\mathbf{c}_p$ and $\mathbf{c}_q$ as $\mathbf{c}_p^c = \mathbf{c}_p - \bar{\mathbf{c}}_p$ and $\mathbf{c}_q^c = \mathbf{c}_q - \bar{\mathbf{c}}_q$ and formulation (4), since $corr(\mathbf{c}_p, \mathbf{c}_q) = corr(\mathbf{c}_q^c, \mathbf{c}_p^c) = cos(\mathbf{c}_p^c, \mathbf{c}_q^c)$; correlation between $\mathbf{e}_p$ and $\mathbf{e}_q$ can be written as:

$$\begin{aligned} corr(\mathbf{e}_p, \mathbf{e}_q) &= corr\big(\ [\mathbf{c}_p^c \cdot \alpha_1',\ \mathbf{c}_p^c \cdot \alpha_2', ..., \mathbf{c}_p^c \cdot \alpha_n'], [\mathbf{c}_q^c \cdot \alpha_1',\ \mathbf{c}_q^c \cdot \alpha_2', ..., \mathbf{c}_q^c \cdot \alpha_n']\big) \\ &= corr\big(\ [\ \|\alpha_1\| cos(\mathbf{c}_p^c, \alpha_1),\ \|\alpha_2\| cos(\mathbf{c}_p^c, \alpha_2), ...,\ \|\alpha_n\| cos(\mathbf{c}_q^c, \alpha_n)\,] \\ &\quad [\ \|\alpha_1\| cos(\mathbf{c}_q^c, \alpha_1),\ \|\alpha_2\| cos(\mathbf{c}_q^c, \alpha_2), ...,\ \|\alpha_n\| cos(\mathbf{c}_q^c, \alpha_n)\,]\ \big) \end{aligned} \tag{5}$$

Where $cos(a, b)$ is the cosine of the angle between the two vectors. As the angle between $\mathbf{c}_p^c$ and $\mathbf{c}_q^c$ gets smaller (equivalent to their correlation getting higher), the difference between $cos(\mathbf{c}_p^c, \alpha_i)$ and $cos(\mathbf{c}_q^c, \alpha_i)$ gets smaller and therefore correlation between $\mathbf{e}_p$ and $\mathbf{e}_q$ gets higher.

Figure 3C shows simulation results for correlation of CT expression profiles versus the observed correlation in the bulk tissue for $1000$ gene pairs.

###### MATLAB code for the simulation

---

```
n = 100
CTCorr = zeros(1, 1000); % correlation of the CT expression profiles
sampleCorr = zeros(1, 1000); % correlation of the gene pair in the bulk tissue
for k = 1:1000
    P = (randn(1,10)*3 + 4);  % generating CT profile for gene P
    Q = (randn(1,10)*3 + 4);  % generating CT profile for gene Q
```

```
    Gs = [P, Q];

    CTCorr(k) = corr(P', Q');

    A = rand(10,  n);
    % normalizing the weight matrix
    for i = 1:n
        A(:, i) = A(:, i) ./(sum(A(:,i)));
    end

    % getting the final vectors
    exps = Gs * A;

    sib = corr(exps');
    sampleCorr(k) = sib(1, 2);
end
```

---

#### 2 Gene variance explained by variance of the marker genes

We examined the results from the Principal Component Regression method from the two sets of marker genes from the same five cell types - Astrocyte, Microglia, Oligodendrocyte, Endothelial and Pyramidal. This was to study the specificity of the model in capturing cellular composition induced variance (refelected specifically in the variation of the marker genes) as oppose to variance induced by a general confounding factor shared among all the genes. The two sets of marker genes are identified independently and have a small overlap (see SFigure 1C and SFigure 2).

#### 3 Figures

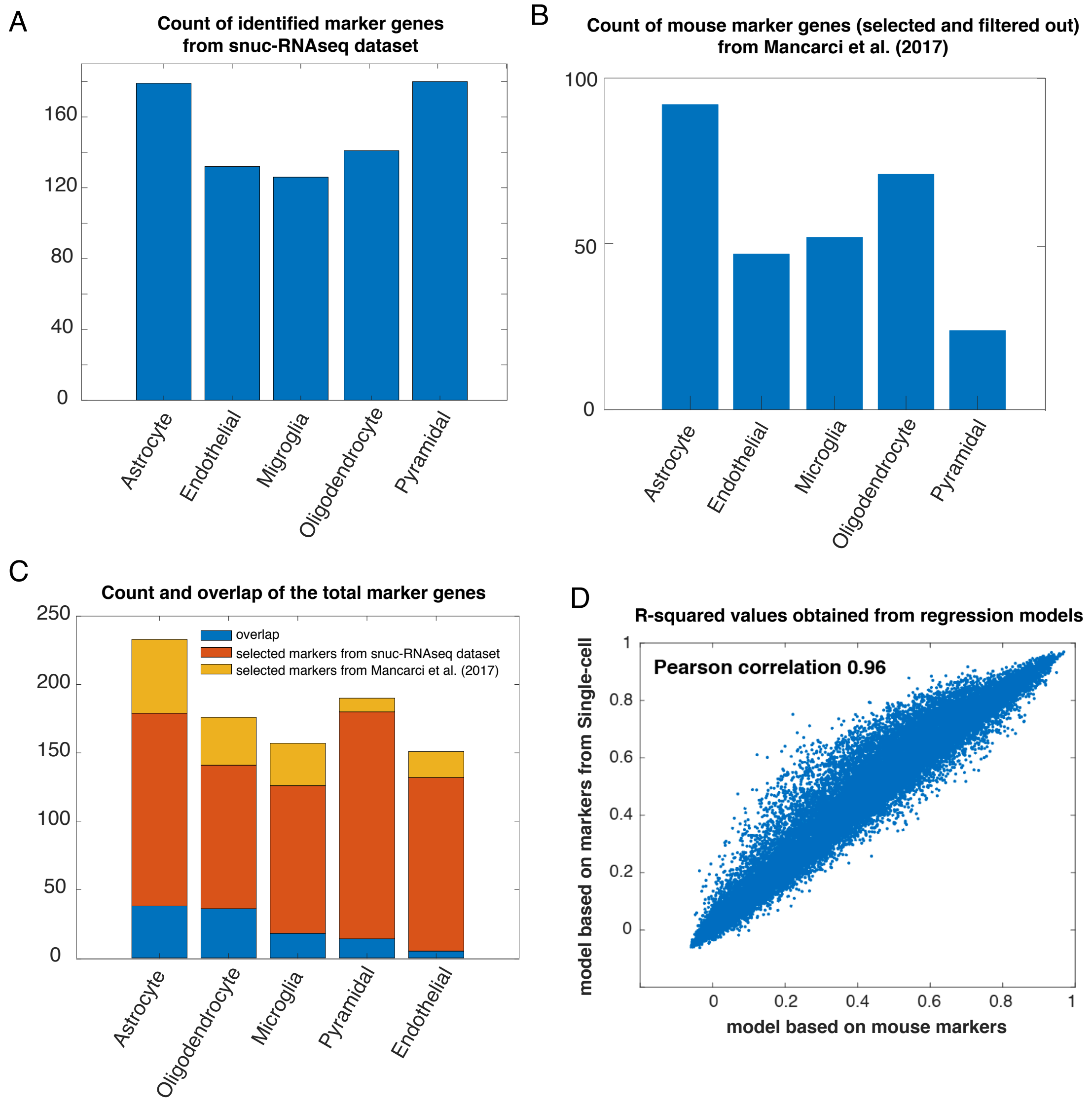


Figure 1: **Comparison of two sets of marker genes.** (A) Count of marker genes identified in snuc-RNAseq dataset. (B) Marker genes from Mancarci et al. (2017). (C) Overlap of the marker genes between the two sources. (D) R-squared values from the two sets of marker genes are highly correlated

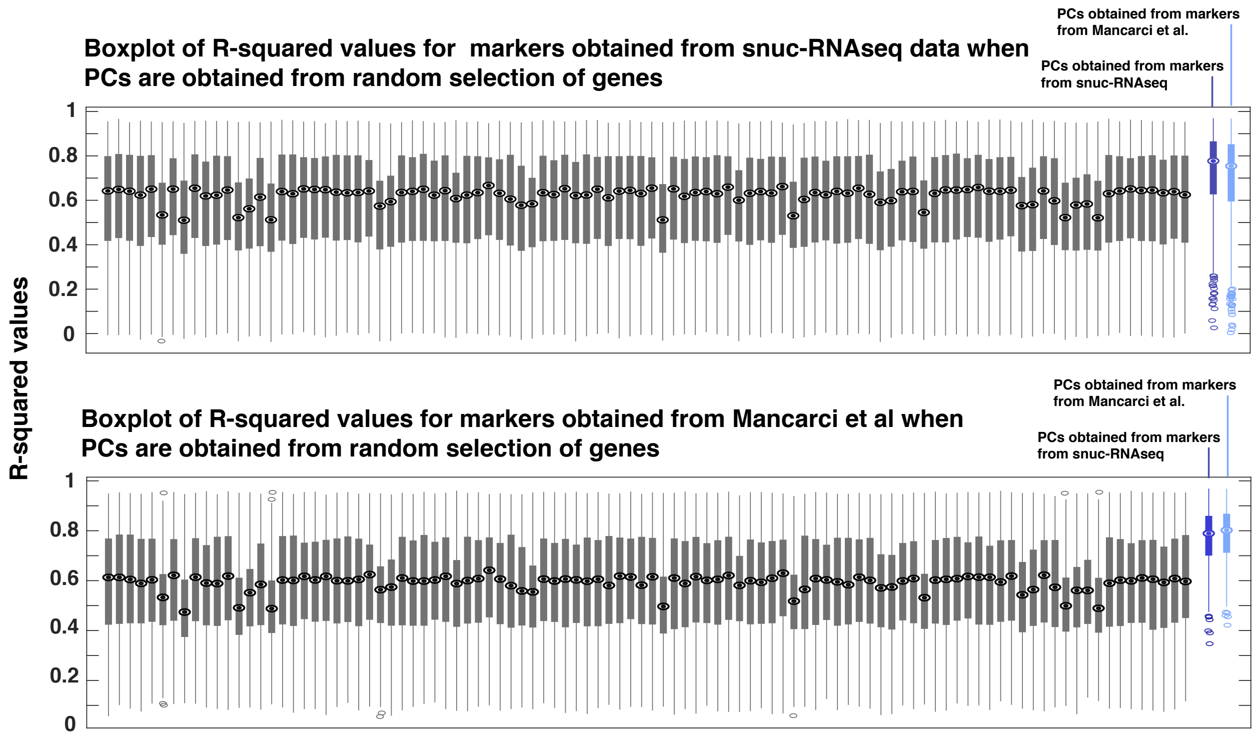


Figure 2: **Comparison of the distribution of $R^2$ values for marker genes, when PCs are obtained from random selection of genes versus when they are obtained from marker genes.**

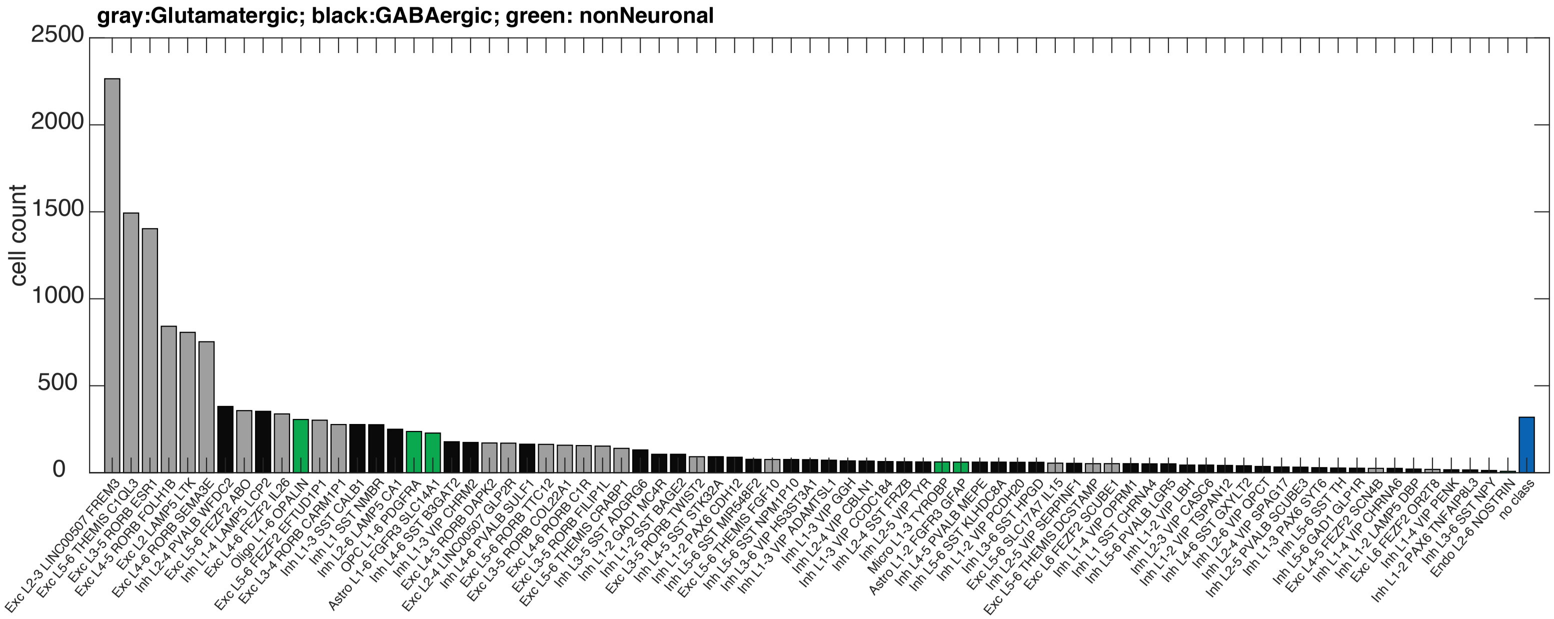


Figure 3: Count of samples (cells) in each cluster of identified cell types for the Allen institude single cell dataset

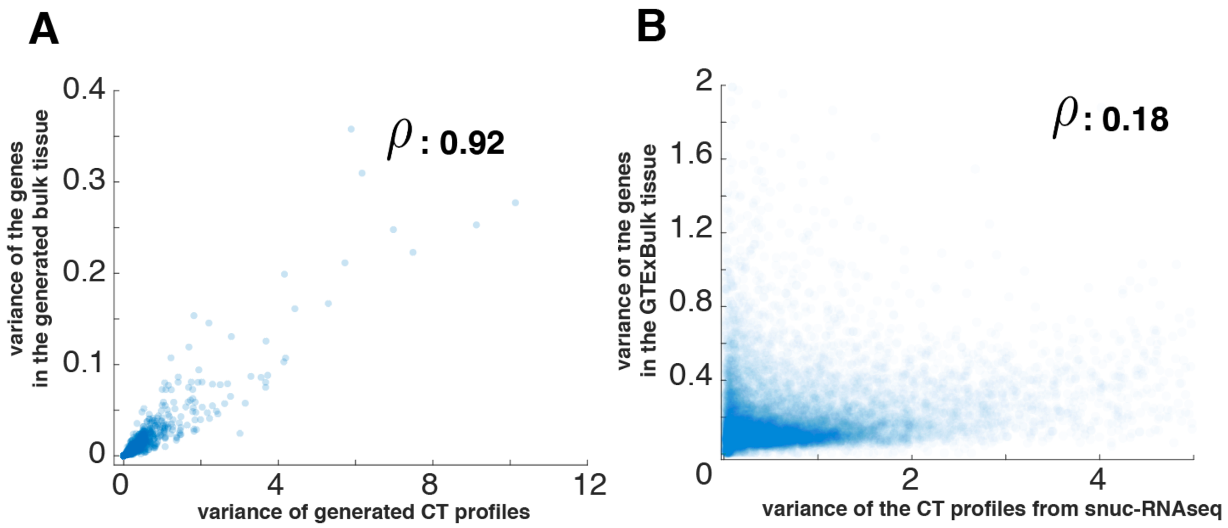


Figure 4: **Variance of the CT profiles versus the variance observed in the bulk tissue.** (A) Results from simulation for $1000$ genes. Each point is data from a gene.(B) Results from data: CT profiles estimated using snuc-RNAseq data, compared to the observed variance in the GTExBulk tissue dataset

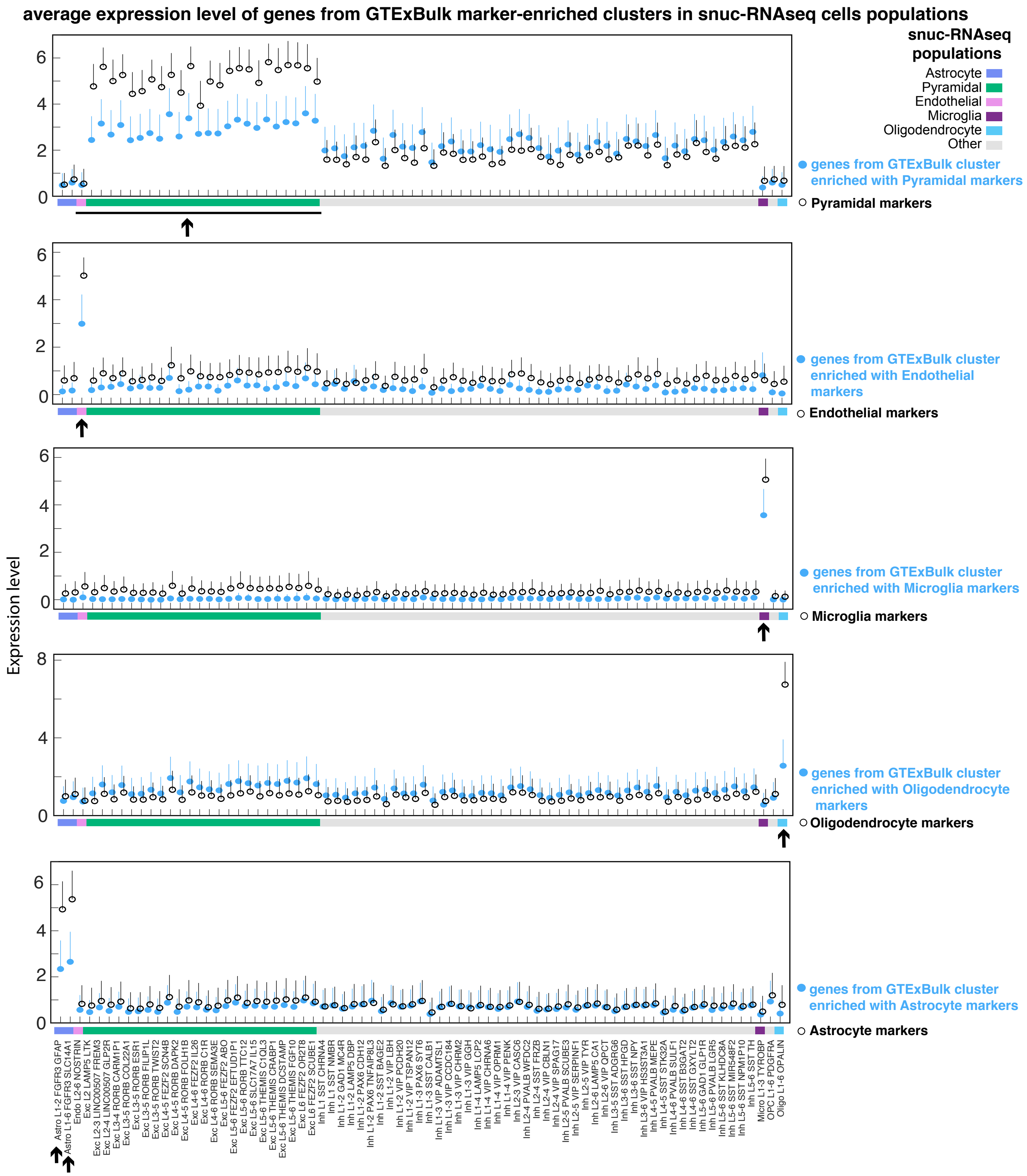


Figure 5: Mean expression level of genes from GTExBulk marker-enriched clusters in different cell populations of the snuc-RNAseq data. Each plot shows the expression levels for genes from GTExBulk clusters enriched with markers of specific cell types (from top to bottom: Pyramidal, Endothelial, Microglia, Oligodendrocyte and Asctrocyte). Lines show the standard deviation. Generally, genes in marker-enriched clusters have higher expression levels in the relevant cell populations -marked by an arrows- compared to other cell populations.

| | GTExBulk | GTEx_residual |
|---|---|---|
| Exc L2-3 LINC00507 FREM3 | 4.304 | 5.53 |
| Exc L5-6 THEMIS C1QL3 | 4.061 | 4.695 |
| Exc L3-5 RORB ESR1 | 3.144 | 3.464 |
| Exc L4-5 RORB FOLH1B | 3.572 | 4.455 |
| Exc L2 LAMP5 LTK | 2.238 | 2.708 |
| Exc L4-6 RORB SEMA3E | 3.01 | 3.332 |
| Inh L2-4 PVALB WFDC2 | 2.45 | 2.704 |
| Exc L5-6 FEZF2 ABO | 2.432 | 2.689 |
| Inh L1-4 LAMP5 LCP2 | 2.385 | 2.594 |
| Exc L4-6 FEZF2 IL26 | 2.518 | 2.656 |
| Exc L5-6 FEZF2 EFTUD1P1 | 2.626 | 2.852 |
| Exc L3-4 RORB CARM1P1 | 2.581 | 2.765 |
| Inh L1-3 SST CALB1 | 2.451 | 2.723 |
| Inh L1 SST NMBR | 2.339 | 2.531 |
| Inh L2-6 LAMP5 CA1 | 2.408 | 2.501 |
| Astro L1-6 FGFR3 SLC14A1 | 2.285 | 2.318 |
| Oligo L1-6 OPALIN | 2.728 | 2.577 |
| OPC L1-6 PDGFRA | 2.326 | 2.429 |
| Inh L4-6 SST B3GAT2 | 2.51 | 2.614 |
| Inh L1-3 VIP CHRM2 | 2.406 | 2.404 |
| Exc L2-4 LINC00507 GLP2R | 2.496 | 2.485 |
| Exc L4-5 RORB DAPK2 | 2.495 | 2.533 |
| Exc L5-6 RORB TTC12 | 2.478 | 2.667 |
| Exc L3-5 RORB COL22A1 | 2.57 | 3.478 |
| Inh L4-6 PVALB SULF1 | 2.437 | 2.555 |
| Exc L4-6 RORB C1R | 2.434 | 2.517 |
| Exc L3-5 RORB FILIP1L | 2.5 | 2.655 |
| Exc L5-6 THEMIS CRABP1 | 2.608 | 3.679 |
| Inh L3-5 SST ADGRG6 | 2.632 | 2.459 |
| Inh L1-2 GAD1 MC4R | 2.493 | 2.454 |
| Inh L1-2 SST BAGE2 | 2.557 | 2.494 |
| Exc L3-5 RORB TWIST2 | 2.624 | 2.561 |
| Inh L4-5 SST STK32A | 2.798 | 2.598 |
| Inh L1-2 PAX6 CDH12 | 2.741 | 2.651 |
| Inh L5-6 SST MIR548F2 | 2.768 | 2.678 |
| Exc L5-6 THEMIS FGF10 | 2.666 | 2.999 |
| Inh L5-6 SST NPM1P10 | 2.902 | 2.749 |
| Inh L3-6 VIP HS3ST3A1 | 2.939 | 2.637 |
| Inh L1-3 VIP ADAMTSL1 | 2.957 | 2.512 |
| Inh L1-3 VIP GGH | 3.12 | 2.952 |
| Inh L2-4 VIP CBLN1 | 3.034 | 2.517 |
| Inh L1-3 VIP CCDC184 | 3 | 2.597 |
| Inh L2-4 SST FRZB | 3.281 | 2.878 |
| Inh L2-5 VIP TYR | 3.11 | 2.749 |
| Inh L4-5 PVALB MEPE | 2.968 | 2.738 |
| Inh L5-6 SST KLHDC8A | 3.188 | 2.559 |
| Inh L3-6 SST HPGD | 3.16 | 2.631 |
| Inh L1-2 VIP PCDH20 | 3.219 | 2.864 |
| Astro L1-2 FGFR3 GFAP | 2.992 | 3.811 |
| Exc L5-6 SLC17A7 IL15 | 3.299 | 2.858 |
| Inh L2-5 VIP SERPINF1 | 3.182 | 2.65 |
| Exc L5-6 THEMIS DCSTAMP | 2.995 | 3.177 |
| Exc L6 FEZF2 SCUBE1 | 4.006 | 3.204 |
| Inh L1-4 VIP OPRM1 | 3.222 | 3.025 |
| Inh L1 SST CHRNA4 | 3.179 | 2.684 |
| Inh L5-6 PVALB LGR5 | 3.198 | 2.596 |
| Inh L1-2 VIP LBH | 3.813 | 3.178 |
| Inh L2-3 VIP CASC6 | 3.472 | 2.896 |
| Inh L1-2 VIP TSPAN12 | 3.337 | 2.878 |
| Inh L4-6 SST GXYLT2 | 3.81 | 2.962 |
| Inh L2-6 VIP QPCT | 3.897 | 3.441 |
| Inh L2-4 VIP SPAG17 | 4.584 | 3.239 |
| Inh L2-5 PVALB SCUBE3 | 4.297 | 3.189 |
| Inh L1-3 PAX6 SYT6 | 4.568 | 3.074 |
| Inh L5-6 SST TH | 5.714 | 3.736 |
| Inh L5-6 GAD1 GLP1R | 5.014 | 3.306 |
| Exc L4-5 FEZF2 SCN4B | 6.177 | 3.819 |
| Inh L1-4 VIP CHRNA6 | 4.883 | 2.977 |
| Inh L1-2 LAMP5 DBP | 6.31 | 2.093 |

Color scale: 2.5 – 6

Figure 6: Ratio of the observed versus the expected link overlap of the links from snuc-RNAseq networks with GTExBulk and GTEx_residual networks. Snuc-RNAseq populations are sorted based on the count of cells, with the top one having the highest count of cells. Generally, the link overlap between GTExBulk and GTEx_residual doesn't change much and there is some level of agreement between snuc-RNAseq population networks and the two bulk networks.

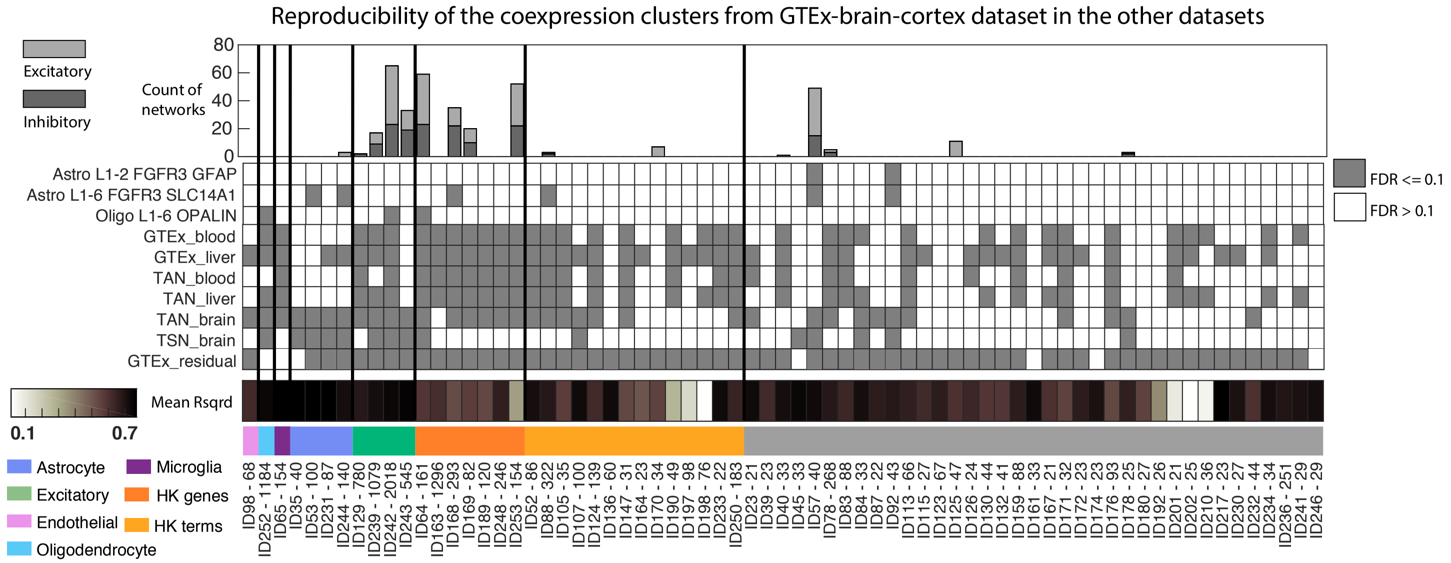


Figure 7: **Reproducibility of clusters identified in GTExBulk in other networks**. Each column in the heatmap (and the same column in the bar plot) shows data for one cluster identified in GTExBulk network. The gray color shows that the cluster had significantly high count of links (FDR ¡= 0.1). The top bar plot shows the count of Excitatory and Inhibitory networks built from populations of cell types identified in snuc-RNAseq data. Rows in the heatmap shows the reproducibility of clusters in different networks. The bottom color bar shows if the cluster has markers of specific cell types, enriched by housekeeping functional terms or genes.

| Clusters | A: Count of links in clusters — GTExBulk | A: GTEx_residual | A: Exc L2-3 LINC00507 FREM3 snuc-RNAseq cell population | B: ratio of the link overlap with the snuc-RNA seq population — GTExBulk | B: GTEx_residual | C: actual link counts retrieved — GTExBulk | C: GTEx_residual | D: GTEx_residual ratio/ GTExBulk ratio |
|---|---|---|---|---|---|---|---|---|
| ID64 - 161 | 3763 | 5504 | 659 | 0.08929 | 0.08358 | 336 | 460 | 0.936 |
| ID78 - 268 | 8497 | 2960 | 155 | 0.008238 | 0.01182 | 70 | 35 | 1.435 |
| ID88 - 322 | 8622 | 2101 | 168 | 0.007307 | 0.01333 | 63 | 28 | 1.824 |
| ID124 - 139 | 1569 | 1767 | 14 | 0.002549 | 0.001698 | 4 | 3 | 0.666 |
| ID129 - 780 | 3.795e+04 | 1.198e+04 | 589 | 0.004505 | 0.006931 | 171 | 83 | 1.538 |
| ID136 - 60 | 506 | 177 | 23 | 0.01581 | 0.02825 | 8 | 5 | 1.787 |
| ID163 - 1296 | 4.894e+04 | 1.09e+05 | 6965 | 0.01645 | 0.0155 | 805 | 1690 | 0.9423 |
| ID168 - 293 | 4455 | 1.439e+04 | 875 | 0.05208 | 0.04129 | 232 | 594 | 0.7928 |
| ID169 - 82 | 547 | 1007 | 18 | 0.01645 | 0.0139 | 9 | 14 | 0.845 |
| ID176 - 93 | 280 | 1335 | 13 | 0.007143 | 0.005993 | 2 | 8 | 0.839 |
| ID189 - 120 | 413 | 2068 | 17 | 0.009685 | 0.003868 | 4 | 8 | 0.3994 |
| ID239 - 1079 | 1.851e+05 | 7.72e+04 | 5085 | 0.01553 | 0.0192 | 2875 | 1482 | 1.236 |
| ID242 - 2018 | 5.319e+05 | 8.314e+04 | 1.124e+04 | 0.01275 | 0.02557 | 6780 | 2126 | 2.006 |
| ID243 - 545 | 3.591e+04 | 2617 | 1178 | 0.01751 | 0.04815 | 629 | 126 | 2.749 |
| ID248 - 246 | 3428 | 2903 | 37 | 0.00175 | 0.001378 | 6 | 4 | 0.7872 |
| ID252 - 1184 | 7.331e+04 | 7141 | 482 | 0.001132 | 0.004481 | 83 | 32 | 3.958 |
| ID253 - 154 | 3101 | 9816 | 1204 | 0.2106 | 0.1201 | 653 | 1179 | 0.5704 |

Networks

Color scale (D): 0 2 4.5

Figure 8: **Representation of the links in a snuc-RNAseq population (Exc L2-3 LinC00507 FREM3). This population was selected because it has the highest count of cells among all the snuc-RNAseq populations and therefore a more complete network regarding the gene count.** **A** shows count of links in each of the GTExBulk clusters (clusters were selected to have 10 or more links in the snuc-RNAseq population). **B** shows ratio of links from GTExBulk and GTEx_Residual networks that overlap with links in the snuc-RNAseq network. **C** shows the actual count of links from snuc-RNAseq network retrieved (has overlap) in GTExBulk and GTEx_residual networks. For the Pyramidal clusters in the green box, although the count of links has decreased, precision has almost doubled for the GTEx_residual network. For the housekeeping cluster in the orange box, count of links retrieved is more than double in GTEx_residual network compared to GTExBulk network and the precision has not changed much.

#### 4 Tables

Table 1: Count of links and genes in each of the networks

| networks | gene count | link count |
|---|---|---|
| GTEx brain cortex | 14,102 | 1,774,291 |
| GTEx blood | 11,348 | 1,312,228 |
| GTEx liver | 16,388 | 1,481,028 |
| TAN brain | 8,761 | 1,118,791 |
| TAN blood | 8,747 | 380,875 |
| TAN liver | 10,227 | 496,476 |
| TSN brain | 6,422 | 358,531 |

| | Count of genes in the networks | Count of inks in the networks |
|---|---|---|
| Exc L2-3 LINC00507 FREM3 | 1.061e+04 | 5.63e+05 |
| Exc L5-6 THEMIS C1QL3 | 1.071e+04 | 5.733e+05 |
| Exc L3-5 RORB ESR1 | 8509 | 3.62e+05 |
| Exc L4-5 RORB FOLH1B | 1.104e+04 | 6.097e+05 |
| Exc L2 LAMP5 LTK | 8530 | 3.638e+05 |
| Exc L4-6 RORB SEMA3E | 9194 | 4.226e+05 |
| Inh L2-4 PVALB WFDC2 | 9107 | 4.109e+05 |
| Exc L5-6 FEZF2 ABO | 1.014e+04 | 5.089e+05 |
| Inh L1-4 LAMP5 LCP2 | 8169 | 3.26e+05 |
| Exc L4-6 FEZF2 IL26 | 9410 | 4.345e+05 |
| Exc L5-6 FEZF2 EFTUD1P1 | 1.081e+04 | 5.73e+05 |
| Exc L3-4 RORB CARM1P1 | 1.034e+04 | 5.153e+05 |
| Inh L1-3 SST CALB1 | 5968 | 1.573e+05 |
| Inh L1 SST NMBR | 7839 | 2.784e+05 |
| Inh L2-6 LAMP5 CA1 | 8656 | 3.421e+05 |
| Astro L1-6 FGFR3 SLC14A1 | 4427 | 7.056e+04 |
| Oligo L1-6 OPALIN | 3286 | 3.234e+04 |
| OPC L1-6 PDGFRA | 3281 | 2.985e+04 |
| Inh L4-6 SST B3GAT2 | 7402 | 1.985e+05 |
| Inh L1-3 VIP CHRM2 | 7583 | 2.017e+05 |
| Exc L2-4 LINC00507 GLP2R | 8894 | 3.172e+05 |
| Exc L4-5 RORB DAPK2 | 8636 | 2.857e+05 |
| Exc L5-6 RORB TTC12 | 1.048e+04 | 4.644e+05 |
| Exc L3-5 RORB COL22A1 | 8420 | 2.521e+05 |
| Inh L4-6 PVALB SULF1 | 8339 | 2.468e+05 |
| Exc L4-6 RORB C1R | 9216 | 3.23e+05 |
| Exc L3-5 RORB FILIP1L | 9276 | 3.215e+05 |
| Exc L5-6 THEMIS CRABP1 | 1.015e+04 | 4.112e+05 |
| Inh L3-5 SST ADGRG6 | 7035 | 1.304e+05 |
| Inh L1-2 GAD1 MC4R | 6328 | 7.98e+04 |
| Inh L1-2 SST BAGE2 | 5777 | 6.454e+04 |
| Exc L3-5 RORB TWIST2 | 7885 | 1.471e+05 |
| Inh L4-5 SST STK32A | 5401 | 5.564e+04 |
| Inh L1-2 PAX6 CDH12 | 7204 | 1.18e+05 |
| Inh L5-6 SST MIR548F2 | 8185 | 1.776e+05 |
| Exc L5-6 THEMIS FGF10 | 9303 | 2.712e+05 |
| Inh L5-6 SST NPM1P10 | 7086 | 1.167e+05 |
| Inh L3-6 VIP HS3ST3A1 | 6367 | 8.461e+04 |
| Inh L1-3 VIP ADAMTSL1 | 5939 | 7.399e+04 |
| Inh L1-3 VIP GGH | 4470 | 4.25e+04 |
| Inh L2-4 VIP CBLN1 | 3773 | 2.924e+04 |
| Inh L1-3 VIP CCDC184 | 5904 | 7.842e+04 |
| Inh L2-4 SST FRZB | 4674 | 4.869e+04 |
| Inh L2-5 VIP TYR | 4724 | 4.851e+04 |
| Inh L4-5 PVALB MEPE | 7049 | 1.278e+05 |
| Inh L5-6 SST KLHDC8A | 5764 | 7.494e+04 |
| Inh L3-6 SST HPGD | 5986 | 8.756e+04 |
| Inh L1-2 VIP PCDH20 | 4353 | 4.205e+04 |
| Astro L1-2 FGFR3 GFAP | 917 | 2964 |
| Exc L5-6 SLC17A7 IL15 | 6698 | 1.194e+05 |
| Inh L2-5 VIP SERPINF1 | 2972 | 2.094e+04 |
| Exc L5-6 THEMIS DCSTAMP | 7535 | 1.737e+05 |
| Exc L6 FEZF2 SCUBE1 | 6841 | 1.388e+05 |
| Inh L1-4 VIP OPRM1 | 2960 | 2.086e+04 |
| Inh L1 SST CHRNA4 | 3136 | 2.304e+04 |
| Inh L5-6 PVALB LGR5 | 3522 | 2.787e+04 |
| Inh L1-2 VIP LBH | 4474 | 5.147e+04 |
| Inh L2-3 VIP CASC6 | 4596 | 5.425e+04 |
| Inh L1-2 VIP TSPAN12 | 2550 | 1.673e+04 |
| Inh L4-6 SST GXYLT2 | 3315 | 2.901e+04 |
| Inh L2-6 VIP QPCT | 1717 | 9126 |
| Inh L2-4 VIP SPAG17 | 1074 | 4576 |
| Inh L2-5 PVALB SCUBE3 | 1516 | 7848 |
| Inh L1-3 PAX6 SYT6 | 2202 | 1.542e+04 |
| Inh L5-6 SST TH | 1450 | 8536 |
| Inh L5-6 GAD1 GLP1R | 856 | 3302 |
| Exc L4-5 FEZF2 SCN4B | 2936 | 3.231e+04 |
| Inh L1-4 VIP CHRNA6 | 409 | 1300 |
| Inh L1-2 LAMP5 DBP | 158 | 304 |

**Table 2** Count of genes (with one or more links) and links in different snuc-RNAseq networks.
